# Nervous system reduction in branched-chain amino acid metabolism disrupts hippocampal neurogenesis and memory

**DOI:** 10.1101/2022.06.10.495711

**Authors:** Khadar Abdi, Ramona M. Rodriguiz, William C. Wetsel, Michelle E. Arlotto, Robert W. McGarrah, Phillip J. White

## Abstract

A role for macronutrient metabolism in learning and memory is supported by numerous epidemiological studies. The *Ppm1k* gene encodes the branched-chain keto acid dehydrogenase (BCKDH) phosphatase that promotes the metabolism of branched-chain amino acids (BCAA). Here we show that nervous system deletion of *Ppm1k* in mice increases BCAA levels in brain tissue but not in plasma. These mice have significant impairments in working memory accompanied by a robust accumulation of DCX+/NeuroD1+ immature neurons within the dentate gyrus granule cell layer. Through single cell RNA sequencing and pathway analysis we identified substantial increases in transit-amplifying cells and immature neurons along with activated hedgehog signaling in *Ppm1k* deficient primary neural stem cells (NSCs). Inhibition of mTOR signaling reversed the effects of *Ppm1k* deletion on neuronal progenitor gene activation in primary NSCs. Together our findings uncover a new molecular link between BCAA metabolism, hippocampal neurogenesis, and cognitive performance.

## INTRODUCTION

Diverse human epidemiological studies and animal models have provided evidence connecting macronutrient metabolism with cognitive performance in both older and younger populations (Dauncey, 2012; Gomez-Pinilla, 2008; Gomez-Pinilla and Tyagi, 2013; Ozawa et al., 2021; Prado and Dewey, 2014). For instance, consumption of high fat diets is correlated with reduced performance on memory tests (Cordner and Tamashiro, 2015; Eskelinen et al., 2008; Greenwood and Winocur, 2005; Vandal et al., 2014). Moreover, high blood glucose, obesity, and type 2 diabetes are each associated with increased risk for development of significant cognitive impairments over time (Biessels and Despa, 2018; Dye et al., 2017). Conversely, intermittent fasting and exercise can improve cognitive performance and delay memory dysfunction (de Cabo and Mattson, 2019; Mandolesi et al., 2018). While studies associating fatty acid and glucose metabolism with learning and memory processes are well-represented, research on how altered amino acid metabolic pathways impact cognition remain very limited. Investigating the effects of dysregulated amino acid metabolism on cognition may provide new insight into the pathophysiology of acquired memory disorders.

Branched-chain amino acids (BCAAs) are a group of essential amino acids (leucine, isoleucine, and valine) that are associated with metabolic syndromes and brain disorders (Arany and Neinast, 2018; Li et al., 2017; Sun et al., 2016; White et al., 2018; White and Newgard, 2019; Xu et al., 2020; Yoneshiro et al., 2019). Beyond their role in protein synthesis, they are metabolized as an energy source, and leucine is further utilized as an essential signaling molecule through the mTOR pathway (Saxton and Sabatini, 2017). BCAAs may also be used as nitrogen donors to produce glutamate and are metabolized into intermediates of the TCA cycle and can serve as paracrine signaling factors that include 3-hydroxyisobutyrate (White et al., 2021). Elevated circulating levels of BCAAs are strongly correlated with type 2 diabetes, insulin resistance, and obesity (Newgard, 2012; White and Newgard, 2019). Serum levels of BCAAs and human mutations in enzymes that regulate their metabolic fate have also been correlated with neurodevelopmental and neurodegenerative disorders (Larsson and Markus, 2017; Trinh et al., 2019; Tynkkynen et al., 2018). Tissue levels of BCAAs are controlled in part by the enzymatic activity of the mitochondrial branched-chain alpha-ketoacid dehydrogenase (BCKDH) complex. This complex is activated through de-phosphorylation by the protein phosphatase M1K (PPM1K), and it is inhibited by the branched-chain *α*-ketoacid dehydrogenase kinase (BDK) (Brosnan and Brosnan, 2006; Li et al., 2017; Lu et al., 2007; Lu et al., 2009). BCKDH is widely expressed in glial cells, neurons, and vascular endothelial cells of the human brain and is predominantly in the active non-phosphorylated state (Bixel et al., 2001; Hull et al., 2018; Suryawan et al., 1998). BDK mutations are associated with autism and epilepsy in both humans and mouse models, while PPM1K mutations are casual for Parkinson’s Disease and are associated with Maple Syrup Urine Disease (Garcia-Cazorla et al., 2014; Novarino et al., 2012; Oyarzabal et al., 2013). While these studies utilizing genetic models that reduce whole body BCAA catabolism describe consequences on heart, liver, and adipose tissue, experiments examining effects of reduced nervous system catabolism of BCAAs are generally lacking.

A major barrier to investigating the role of nervous system BCAA catabolism involves the early postnatal lethality after eliminating BCAA catabolism in the whole animal through BCKDH deletion (Homanics et al., 2006). Additionally, viable animal models with whole body reductions in BCAA catabolism, such as through *Ppm1k* deletion, significantly increase circulating levels of BCAAs to neurotoxic levels, thereby complicating interpretation of nervous system-specific roles for the pathway (Lu et al., 2009). To circumvent these limitations and functionally test the role of BCAA catabolism in the nervous system, we generated conditional knockout mice in which we deleted the BCKDH phosphatase, *Ppm1k*, from the nervous system at birth. This approach allowed us to reduce, rather than eliminate, BCAA catabolism in the nervous system without increasing circulating levels of BCAAs.

Mouse behavioral testing revealed significant negative consequences of *Ppm1k* deletion on spatial working memory. Because spatial memory can be regulated by hippocampal neurogenesis, a process that is metabolically demanding, we further employed immunohistochemical analysis of mouse brains and single cell RNA sequencing of neural stem cells. Our results reveal a previously undescribed role for BCAA metabolism in memory processes, hippocampal neurogenesis, and neuronal differentiation.

## RESULTS

### Behavioral screen of mice with *Ppm1k* nervous system deletion identifies a role in memory

To evaluate the consequences of reduced BCAA catabolism in the nervous system, we crossed *Ppm1k*^flox/flox^ mice with the Nestin-Cre driver to generate Nestin-cre; *Ppm1k* ^Flox/Flox^ conditional knockout mice (cKO) harboring nervous system-specific deletion of the *Ppm1k* gene (**Figure 1A**). Successful deletion of *Ppm1k* was verified by qPCR and western blots showing nearly complete loss of *Ppm1k* mRNA and protein compared to *Ppm1k*^Flox/Flox^ control mice (Ctrl) (**Figure 1B**). *Ppm1k* cKO mice were fertile, viable, and showed no discernible signs of developmental or growth abnormalities.

**Figure 1.**
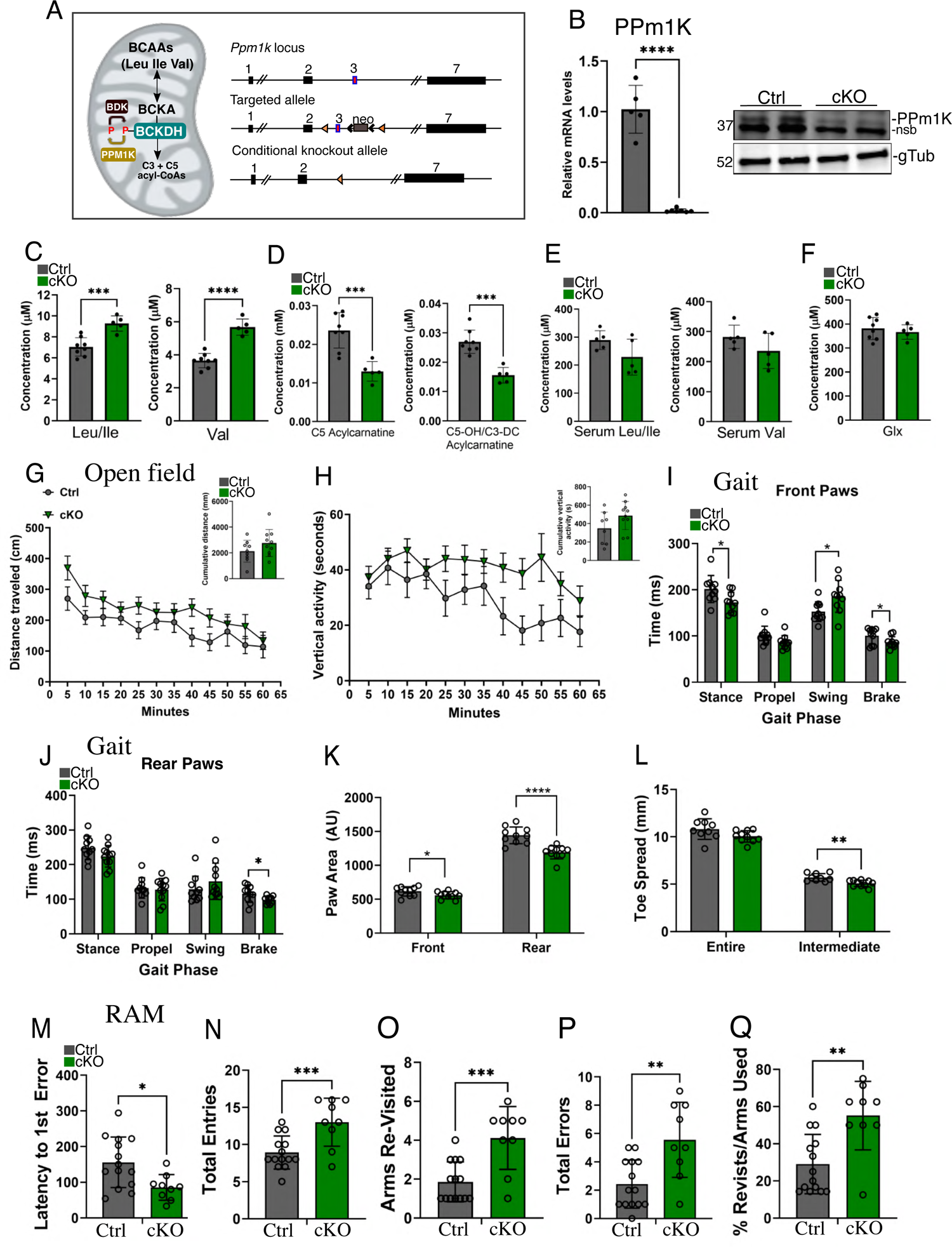
Nervous system deletion of *Ppm1k* results in impaired working memory. A) Illustration of the BCAA catabolic pathway (left) and the mouse *Ppm1k* gene locus (right) showing exon 3 targeted with loxp sites creating the conditional deletion. B) QPCR and western blots from control and *Ppm1k* cKO whole brain samples. Ctrl n = 5, cKO n = 8; ****p < 0.0001, Student’s t-test. C) Quantification of the absolute concentration of branched-chain amino acids from control and ***Ppm1k*** cKO brain tissue. Ctrl n = 8, cKO n = 5; ***p < 0.001, ****p < 0.0001, Student’s t-test. D) Quantification of the absolute concentration of C5 and C5-OH/C3-DC acylcarnitines in brain samples from control and ***Ppm1k*** cKO mice. Ctrl n = 8, cKO, n = 5; ***p < 0.001, Student’s t-test. E) Quantification of the concentrations of BCAAs in serum from control and ***Ppm1k*** cKO mice. Not significant; Ctrl n = 5, cKO n = 5. F) Quantification of the absolute concentration of glutamine/glutamic acid (Glx) from control and PPM1K cKO brain tissue. Not significant; Ctrl n = 8, cKO n = 5. G) Locomotor activity (distance traveled in cm) in 5-min segments over 60 min in the open field. *Inset:* cumulative distance traveled; there was a tendency for locomotion to be higher in the cKO than in the Ctrl mice. Ctrl n = 9, cKO n =10; p=0.083, Ctrl vs. cKO (Student’s t-test). H) Vertical activity (beam-breaks) in 5-min segments over 60 min in the open field. *Inset*: cumulative vertical activity (*p < 0.05, Student’s t-test). Ctrl n = 9, cKO n =10; **p <0.01, Ctrl vs. cKO (repeated measures ANOVA). I) Time (ms) spent in each of the four phases of gait with the fore limbs. Ctrl n = 10, cKO n =10; *p < 0.05, repeated measures ANOVA. J) Time (ms) spent in each of the four phases of gait with the hind limbs. Ctrl n = 10, cKO n =10; *p < 0.05, repeated measures ANOVA. K) Fore paw and hind paw areas in the gait test. Ctrl n = 10, cKO n =10; *p < 0.05, ***p < 0.001, multivariate ANOVA. L) Toe spreads and inter-toe distance in the gait test. Ctrl n = 10, cKO n = 10; ***p < 0.001, multivariate ANOVA. M) Latency to commit the first error in the radial maze. Ctrl n = 14, cKO n =9; **p < 0.05, independent samples two-tailed Student’s t-test. N) Total number of entries into the arms of the radial maze. Ctrl n = 14, cKO n = 9; ***p < 0.001, independent samples two-tailed Student’s t-test. O) Total numbers of arms revisited in the radial maze. Ctrl n=14, cKO n=9; ***p < 0.001, independent samples two-tailed Student’s t-test. P) Total numbers of errors committed in the radial maze. Ctrl n=14, cKO n=9; **p < 0.01, independent samples two-tailed Student’s t-test. Q) Percent errors relative to entries in the radial maze. Ctrl n =14, cKO n = 9; **p < 0.01, independent samples two-tailed Student’s t-test.

To evaluate if *Ppm1k* cKO mice had the expected disruption in BCAA catabolism, we measured amino acids and acylcarnitines in brain samples from control and *Ppm1k* cKO mice, which demonstrated that BCAAs were selectively elevated in *Ppm1k* cKO brain tissue while other amino acids remained at equivalent levels to controls (**Figure 1C and S1**). Consistent with a defect in BCAA catabolism, levels of the C3 and C3-OH/C5-OH acylcarnitines, both downstream metabolites of BCAA oxidation, were reduced in *Ppm1k* cKO brains (**Figure 1D**). Serum levels of BCAAs were similar between *Ppm1k* cKO and control mice, consistent with a brain-specific loss of *Ppm1k* gene expression (**Figure 1E**). Importantly, neural deletion of *Ppm1k* did not alter abundance of glutamate/glutamine in the brain (**Figure 1F**). Likewise, levels of the aromatic amino acids, tyrosine and phenylalanine, precursors for dopamine and norepinephrine, which compete with BCAA for transport across the blood brain barrier via the large neutral amino acid transporter-1 (LAT-1) (Coppola et al., 2013), were equivalent between *Ppm1k* cKO and controls, suggesting their transport into brain tissue was undiminished (**Figure S1**).

We conducted a battery of behavioral tests on 10-12-month-old male and female *Ppm1k* mice to evaluate their motor and memory functions (all statistics are displayed in **Suppl. Table S1 & S2** and Figure legends). Motor performance was examined in the open field (Seibenhener and Wooten, 2015). No genotype effects were observed for locomotor activity when the data were analyzed at 5-min intervals over the 60-min test (**Figure 1G**). Collapsing the data as cumulative distance found a trend for locomotion to be higher in the mutants than control mice (**Figure 1G**, *inset*). By comparison, rearing activity was significantly higher in the mutants than the controls according to both methods of analyses (**Figure 1H**). Hence, *Ppm1k* cKO mice appear to be mildly hyperactive relative to the *Ppm1k*^Flox/Flox^ controls.

Besides examining motor performance, anxiety-like behaviors were also evaluated in the open field. Distance traveled and time spent in the center zone failed to reveal any significant differences between genotype (**Figures S2A and S2B**). Thus, *Ppm1k* cKO mice do not display anxiety-like responses.

In a further examination of motor performance, we evaluated several different parameters for gait. An analysis of the phases of gait with the fore paws determined that stance and brake were significantly lower in cKO mice, while swing was higher than in controls, and no genotype difference was noted for propel (**Figure 1I**). By contrast, the phases of gait by the rear paws were similar between the genotypes, except for brake, which was shorter (**Figure 1J**). In addition, stride lengths for the front and rear paws of the different genotypes were almost superimposed, as were the rear stride width, front- and hind-limb gait angles, homolateral, homologous, and contralateral couplings, and running speeds (**Figures S2C-G**). In comparison, the fore-paw width was narrower in the cKO than Ctrl mice, and paw areas were reduced which appeared due to the diminished toe spreads (toe 2 to toe 4) in the mutant animals (**Figures 1K and 1L and S2C**). Taken together, motor performance in cKO mice appears to be mildly affected relative to control animals.

Since levels of BCAAs have been correlated with brain disorders in humans, we examined spontaneous working memory using the elevated 8-arm radial maze. cKO mice were deficient on all indices of testing in the maze (**Figures 1M-Q and S2H-K**). For instance, the mutants had a shorter latency to enter the first arm more readily (**Figure 1M**), made more entries into the arms (**Figure 1N**), revisited more arms (**Figure 1O**), made more total errors (**Figure 1P**), committed a higher percentage of errors relative to the numbers of arm entries (**Figure 1Q**), tended to use more arms overall (**Figure S2H**), made more perseverative errors (**Figure S2I**), and engaged in more arm entries after the first error to repeat (**Figure S2J**). The numbers of initial arm entries to repeat were not different between the genotypes (**Figure S2K**). Since the open field results for anxiety-like behavior failed to detect any significant genotype effects that could have confounded the radial maze results, the present findings suggest working memory is disturbed in cKO mice.

### PPM1K deletion promotes accumulation of immature neurons in the dentate gyrus

Gross immunohistochemical analysis of coronal sections of whole mouse brains that were stained with cresyl violet showed no obvious defects in brain anatomy (**Figure S3A**). Cortical thickness and layering, formation of the corpus collosum and other brain regions all appeared equivalent between genotypes. Additionally, we did not observe any brain phenotypes related to classical neurodegenerative indices such as increases in the levels of phospho-Tau in *Ppm1k* cKO mice at 12 months of age (**Figure S3B**). Given the hippocampus’s essential role in memory processes and *in situ* mRNA expression data from the Allen Brain bank showing high BCKDH and PPM1K expression in hippocampus (**Figure S3C**), we next focused our analysis on this brain region. To evaluate whether the morphology of the hippocampal formation remained intact, we stained hippocampi using NeuN, a marker for mature neurons. We observed normal morphology and the neuronal densities in hippocampal CA1, CA3, and dentate gyrus were unchanged suggesting their overall organization remained intact after deletion of *Ppm1k* (**Figure S3D**).

The hippocampus is one of two brain regions containing adult neural stem cells capable of generating newborn neurons during postnatal development and throughout adulthood in mammals (Eriksson et al., 1998; Kempermann et al., 2018; Moreno-Jimenez et al., 2019; Toda et al., 2019). Neuronal progenitors born in the subgranular zone (SGZ) migrate locally into the granular cell layer (GCL) to integrate with other cells and to differentiate into mature neurons. Mature neurons send their dendrites to the molecular layer (ML) and the axons from these neurons project through the mossy fibers to synapse on to the CA3 region to receive input from the entorhinal cortex. This tri-synaptic circuit is required for spatial pattern separation, and neurogenesis is directly implicated in spatial memory (Clelland et al., 2009). Labeling of control and *Ppm1k* cKO mouse hippocampus with the newborn neuron marker doublecortin, DCX, revealed a nearly10-fold increase in immature neurons within the granular cell layer (GCL) as well as increased numbers of DCX+ cells within the SGZ (**Figure 2A**). Many of the immature neurons embedded within the GCL in the cKO brains possessed elaborate dendrites and spines but retained strong DCX label throughout. The DCX+ immature neuronal cells were not observed anywhere else in the surrounding tissue, nor could they be found outside any known neurogenic zones in the mouse brain. Interestingly, many DCX+ neurons located in the GCL in cKOs were also positive for the mature neuron marker, NeuN, though their intensity of NeuN staining appeared weaker (**Figure 2B**). We also labeled hippocampal sections with NeuroD1, an early inducer of the neuronal fate that retains expression in newborn neurons prior to their final maturation stage, then becomes downregulated in fully mature neurons (Gao et al., 2009). Consistent with our observation with DCX, NeuroD1+ cells were abundant in the GCL of *Ppm1k* cKO mice compared to control mice, confirming their immature characteristics (**Figure 2C**). Importantly, we determined that Cre expression from the Nestin-cre driver is not responsible for this dentate gyrus phenotype, as Nestin-Cre;*Ppm1k*^WT^ contain similar numbers of DCX+ cells in both their SGZ and GCL compared to *Ppm1k*^flox/flox^ controls (**Figure S3E**).

**Figure 2.**
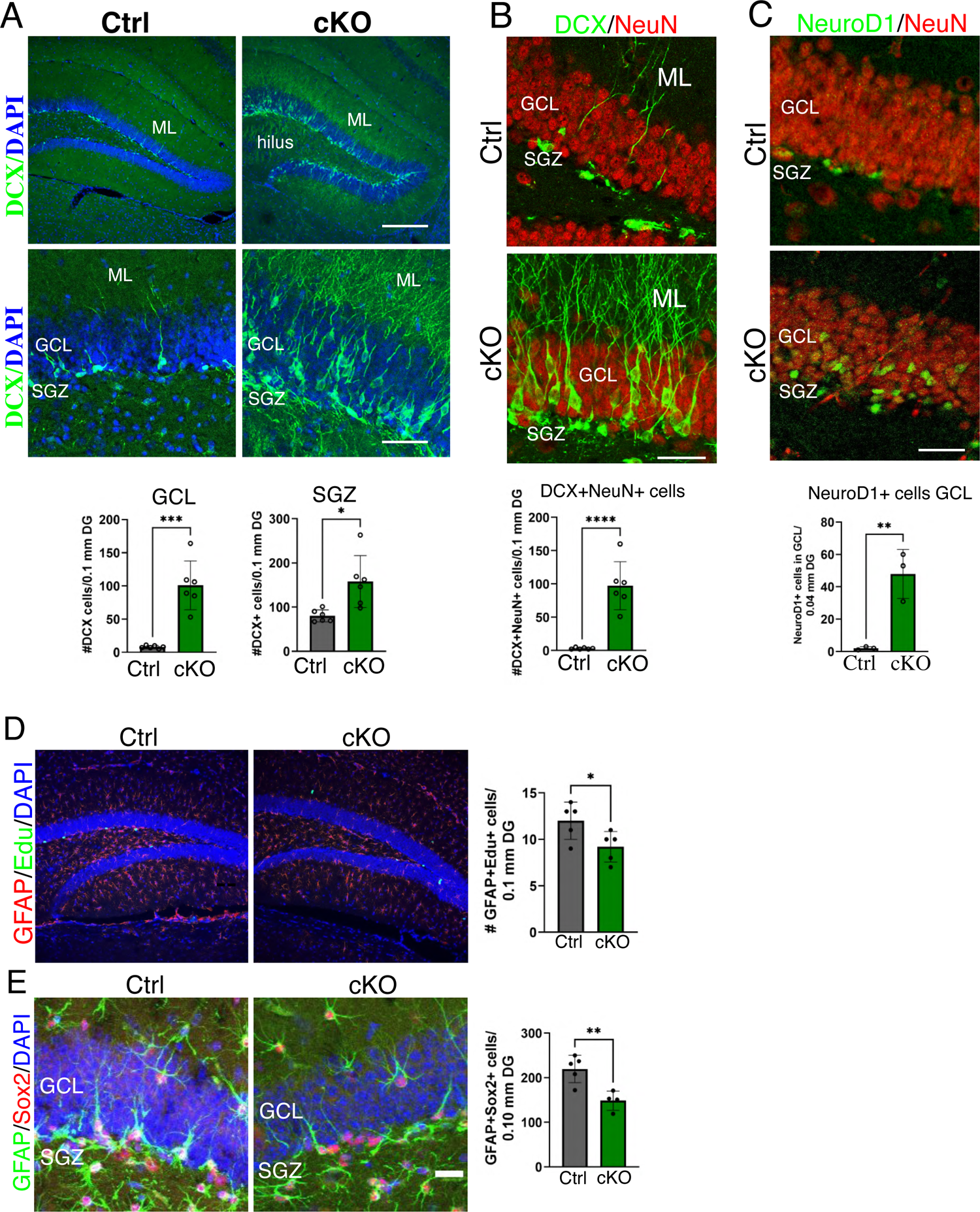
*Ppm1k* cKO mice accumulate immature neurons in the granular cell layer and subgranular zone of the dentate gyrus. A) Representative wide field and higher magnification images from 6-month-old control and *Ppm1k* cKO mouse hippocampal sections stained for doublecortin (DCX) in green and DAPI in blue. Below: Quantification of the number of DCX+ cells found in the GCL of control and *Ppm1k* cKO mice per 0.1 mM of coronal tissue of the dentate gyrus. Ctrl n = 6, cKO n = 6; ***p < 0.001. Quantification of the number of DCX+ cells found in the SGZ of control and *Ppm1k* cKO mice per 0.1 mM of coronal tissue of the dentate gyrus. Ctrl n = 6, cKO n = 6; *p < 0.05. Subgranular zone (SGZ), granular cell layer (GCL), molecular layer (ML). Scale bars = 150 μm, and 40 μm. B) Representative images from 6-month-old control and *Ppm1k* cKO mouse coronal hippocampal sections stained for DCX in green and NeuN in red. Below: Quantification of the number of cells positive for both DCX and NeuN in the GCL of control and *Ppm1k* cKO mice per 0.1 mM of coronal tissue of the dentate gyrus. Ctrl n = 6, cKO n = 6; ***p < 0.001. C) Representative images from 6-month-old control and *Ppm1k* cKO mouse coronal hippocampal sections stained DCX in green and NeuroD1 in red. Below: Quantification of the number of cells positive for NeuroD1 in the GCL of control and *Ppm1k* cKO mice per 0.04 mM of coronal tissue of the dentate gyrus. Ctrl n = 3, cKO n = 3; ***p < 0.001. D) Representative images from 6-month-old control and *Ppm1k* cKO mouse coronal hippocampal sections stained for GFAP in red, Edu in green, and DAPI in blue. Right: Quantification of the number of neural stem cells positive for both GFAP and Edu in the SGZ of control and *Ppm1k* cKO mice per 0.1 mM of coronal tissue of the dentate gyrus. Ctrl n = 5, cKO n = 5; *p < 0.05. Scale bars = 150 μm. E) Representative images from 6-month-old control and *Ppm1k* cKO mouse coronal hippocampal sections stained for GFAP in green, Sox2 in red, DAPI in blue. Scale bars = 150 μm. Right: Quantification of the number of cells that are GFAP and Sox2 double positive in the SGZ of control and *Ppm1k* cKO mice per 0.1 mM of coronal tissue of the dentate gyrus. Ctrl n = 5, cKO n = 5; **p < 0.01. All p values were determined by independent samples two-tailed Student’s t-test

To determine whether increased proliferation of neural stems cells was responsible for the higher numbers of immature neurons in the dentate gyrus, we quantified dividing neural progenitor cells within the SGZ, identified by 5-Ethynyl-2′-deoxyuridine (EdU) GFAP co-labeling. In contrast to the DCX staining, 6-month-old *Ppm1k* cKO brains displayed a modest reduction in Edu/GFAP positive cells along the SGZ (**Figure 2D**). Since this reduced proliferation could be reflective of a reduction in the neural progenitor pool, we quantified the number of adult progenitors within the SGZ using co-labeling of Sox2 and GFAP. We observed that *Ppm1k* cKO mice had fewer GFAP/Sox2 double positive cells than controls, pointing to a loss in the number of adult neural stem cells (**Figure 2E**).

### Loss of PPM1K induces neural stem cell activation and differentiation

To evaluate how PPM1K may regulate adult neural stem cell (NSC) activity, differentiation, and cellular transition states we conducted single cell RNA sequencing analysis in adult NSCs cultured in maintenance media following *Ppm1k* deletion. Adult NSCs lacking PPM1K were produced by adding adenovirus expressing Cre-IRES-GFP to *Ppm1k*^flox/flox^ cultures while control cells were generated by adding the same adenovirus to *Ppm1k*^wt/wt^ cultures. After 68 hours, cells were sorted for GFP fluorescence to exclude non-transduced cells (**Figure 3A**). We performed single cell RNA sequencing on *Ppm1k* KO and control adult NSC cultures using 10,000 cells for each condition. The initial unsupervised UMAP profile contained 16 total clusters (**Figure S4A**). Since these initial 16 clusters exceed the number of known individual cell types generated by NSCs, we reasoned that some clusters likely represented states within the same cell type. To refine the initial 16 clusters in our UMAP into known cell types and cellular states derived from adult NSCs, we used markers defining NSC stages and their progeny described in recent publications (**Figures 3B and 3C**) (Basak et al., 2018; Dulken et al., 2017). Four major subgroups were identified using the markers representing undifferentiated NSCs (udNSCs, using quiescent markers), activated NSCs (aNSCs), transit amplifying cells (TA), and an immature neuronal population (iNeu) (**Figure 3D**). We then compared the refined UMAPs of Ctrl and *Ppm1k* KO cultures to identify distinct differences in their profiles (**Figure 3E**). Control cultures had a UMAP profile with a denser large central cluster representing udNSCs, whereas *Ppm1k* KO cultures were represented by a large shift away from the central cluster and towards aNSCs and differentiation into TA stages (**Figure 3E**). Principal component analysis (PCA) confirmed a clear lineage trajectory away from the udNSC state in cells with *Ppm1k* KO along with an increased number of cells containing markers for TA cells and immature neurons (**Figure 3F**). *Ppm1k* KO cultures had more aNSCs, TA, and iNeu cells than controls, revealing an overall transition into activation and differentiation in the absence of PPM1K (**Figure 3G**). These results demonstrate that PPM1K plays an important role in the regulation of adult NSC activation and differentiation, and its loss promotes NSC activation and lineage trajectory into neuronal progenitors.

**Figure 3.**
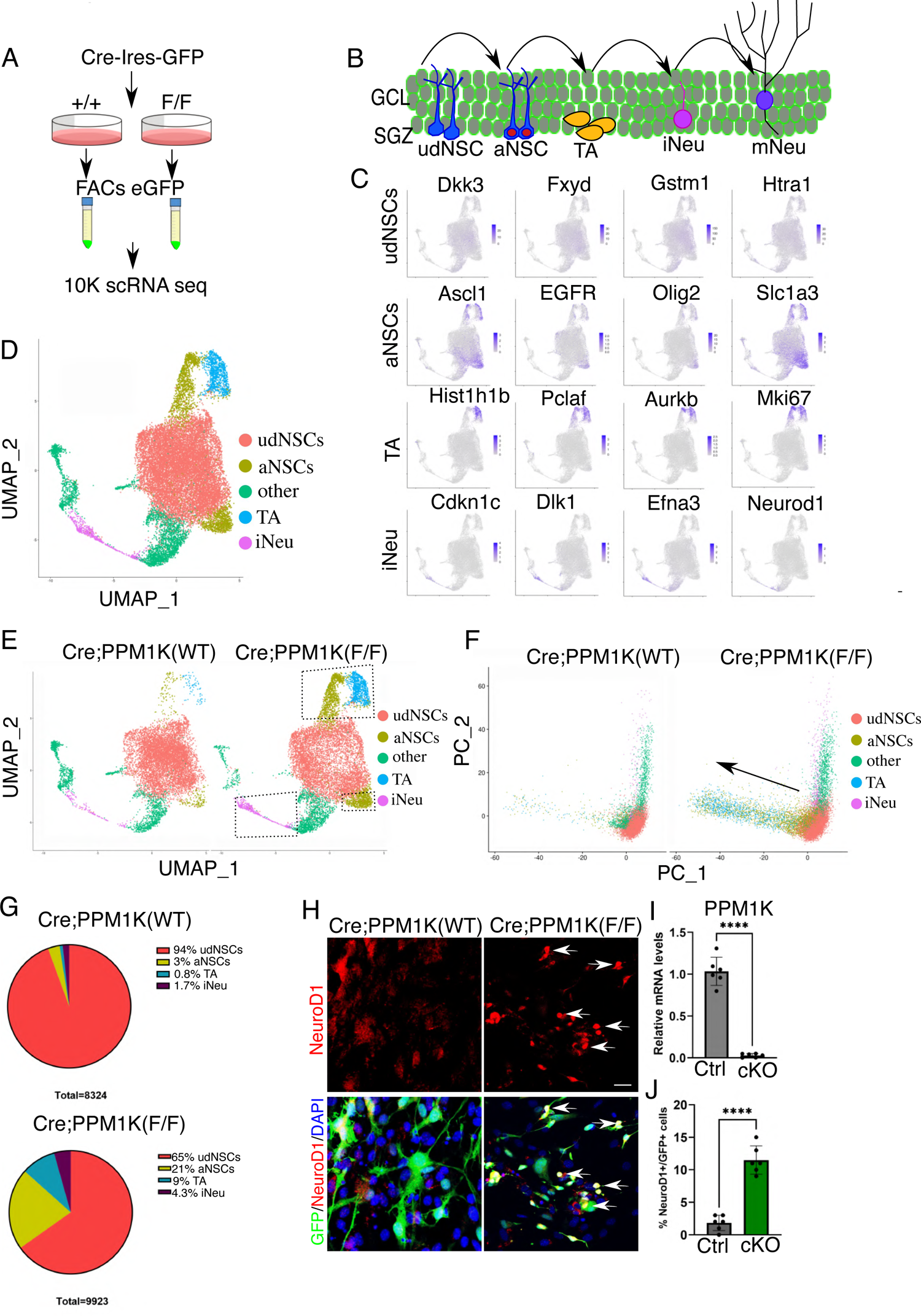
Single cell RNA sequence reveals enhanced NSC activation and differentiation in *Ppm1k* KOs. A) Schematic illustrating the method used to perform scRNA sequencing. B) Illustration of the lineage stages of adult neural stem cells (NSCs) differentiation into mature neurons within the dentate gyrus; undifferentiated neural stem cells (udNSCs), activated neural stem cells (aNSCs), transit amplifying cells (TA), immature neuron (iNeu), and mature neuron (mNeu). C) Gene markers for specific neural stem cell subtypes plotted on the UMAP. Four cell type specific markers are show for each udNSCs, aNSCs, TA, and iNeu populations. D) Refined UMAP showing the four major cell types identified through marker selection. The cell population marked “other” was not identifiable by cell subtype markers. E) Comparison of refined UMAPs from control and *Ppm1k* KO NSC. Color is coded similar to (D). Dashed boxes outline populations increased in the knockout condition. F) PCA plots generated from refined cell subtype maps of control and PPM1K KO NSCs. Color is coded similar to (E). Arrow signifies the increase in activated NSCs and TA populations in the knockout condition. G) Pie graph showing the percentages of each cell subtype present in control and *Ppm1k* KO NSC cultures. H) Representative confocal images from control and *Ppm1k* KO NSCs labeled for GFP (green), NeuroD1 (red), and DAPI (blue). Arrows point to cells that are both positive for GFP and NeuroD1. Scale bar = 20 μm. I) Quantification of the relative mRNA expression levels for *Ppm1k* in control and *Ppm1k* KO cultures. ****p < 0.0001; Ctrl n = 6, cKO n = 6. All data are presented as mean +/- SD. J) Quantification of the percentage of iNeu cells that are positive for both GFP and NeuroD1 in control and *Ppm1k* KO cultures. Ctrl n = 6, cKO n = 6; *p < 0.001. All p values were determined by independent samples two-tailed Student’s t-test

The presence of increased numbers of immature neurons in *Ppm1k* cKO cultures is consistent with our observations in *Ppm1k* cKO brains where we observed an accumulation of immature neurons within the GCL. To determine whether we could visually identify cells with immature neuronal markers in NSC cultures, we performed confocal microscopy with both control and *Ppm1k* cKO neural stem cell cultures expressing Cre-IRES-GFP. After 3 days in growth conditions, we observed the appearance of GFP+ cells that were NeuroD1 positive in *Ppm1k* cKO cultures that were rarely seen in control cultures (**Figures 3H-J**). These results point to a consistent increase in NSC activation and differentiation in *Ppm1k* KO cultures and could represent altered quiescence or fate specification pathways during loss of PPM1K protein. It is noteworthy that loss of PPM1K is sufficient to elicit these effects in the absence of any changes in growth factors or nutrient conditions typically required to initiate differentiation.

### GSEA pathway analysis identifies hedgehog signaling activation in *Ppm1k* KO NSCs

We next sought to uncover mechanisms that may be driving *Ppm1k* deleted NSCs into the activated and differentiated neural progenitors seen in the single cell analysis of *Ppm1k* KOs. We first performed differential expression (DE) analysis between control and KO cells in the undifferentiated NSC cluster (**Figure 4A**). DE analysis identified 311 upregulated genes and 228 downregulated genes meeting a cut-off of p < 0.05 and Log2 fold-change of +/- 0.25. Differentially upregulated genes between control and *Ppm1k* KOs represented a class of genes involved in proliferation and early markers for activated NSC states such as fatty acid binding protein 7 (Fabp7), cyclin D1 (Ccnd1), and pleiotrophin (Ptn) (**Figure 4B**). Differentially downregulated genes included markers of quiescent NSCs such as Apoe and Cst3, consistent with the observed reduction in the udNSC population in *Ppm1k* KOs (**Figure 4B**) (Basak et al., 2018).

**Figure 4.**
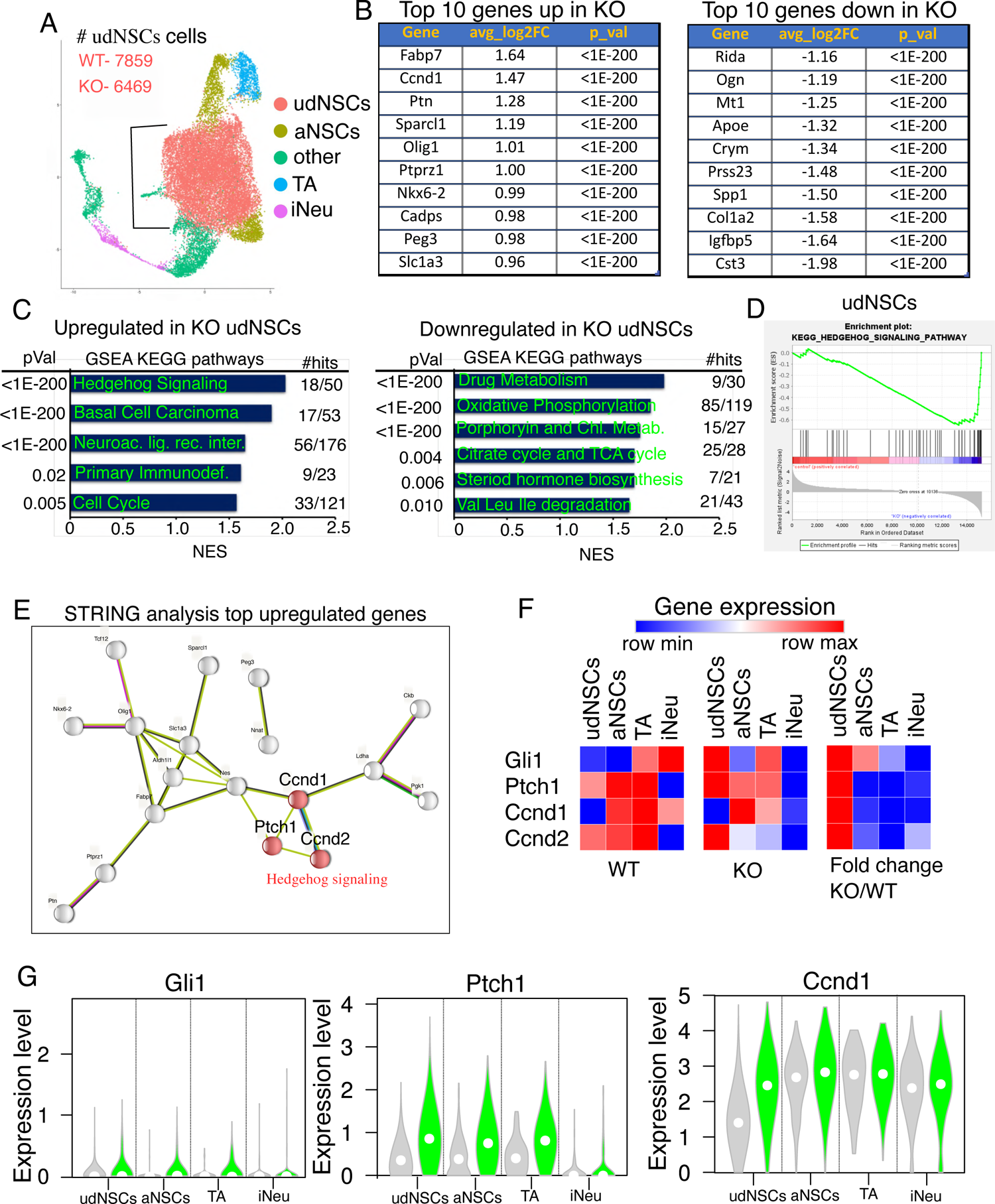
Gene set enrichment analysis uncovers upregulated hedgehog signaling in *Ppm1k* Kos. A) The refined UMAP showing all four major cell types identified and with the udNSC population cell numbers noted. B) Tables listing top 10 upregulated and downregulated genes in the udNSC population from scRNA sequencing. Average Log2 fold change and p values are presented for each gene. C) Gene set enrichment analysis (GSEA) showing normalized enrichment scores and p values of upregulated and downregulated KEGG pathways that are differentially expressed between control and *Ppm1k* KO NSCs in the udNSC population. For all pathways FDR < 0.1, *p < 0.05. D) Gene set enrichment plot for the hedgehog signaling pathway in the udNSC population. E) STRING protein analysis of the top 30 upregulated differentially expressed genes (*Ppm1k* KO NSCs compared to control NSCs) from the udNSC group with unconnected nodes removed. Top three genes represented in the hedgehog signaling pathway are colored. F) Heatmaps of top gene hits in the hedgehog signaling pathway identified through GSEA. Gene expression levels were followed along the NSC lineage trajectory path to produce maps showing lowest to highest expression over cell types. First two heatmaps show average expression from control samples and *Ppm1k* KO samples across cell types. Third heatmap shows the differential expression levels between control and *Ppm1k* KO cells along each cell type. G) Violin plots are shown for three hedgehog signaling related genes identified through pathway analysis.

To identify novel pathways that could induce activation toward neuronal differentiation connected to the *Ppm1k* cKO phenotype in NSCs, we performed gene set enrichment analysis (GSEA) on the udNSC population data. Using the KEGG database, we identified hedgehog signaling as the top upregulated pathway, with a 2.03 NES score (**Figures 4C and 4D**). Additional upregulated pathways, such as basal cell carcinoma, encompassed many of the same hedgehog signaling pathway and cell cycle genes (**Table S3**). Drug metabolism and oxidative phosphorylation were the top two downregulated pathways in the *Ppm1k* cKO condition, with NES scores of 1.98 and 1.85, respectively (**Figure 4C**). As expected, the sixth downregulated pathway identified in the GSEA included degradation of the branched chain amino acids, leucine, isoleucine, and valine, consistent with PPM1K’s role BCAA catabolism (**Figure 4C**). Additional reduced pathways on the list that shared top NES scores included steroid hormone biosynthesis and the TCA cycle (**Figure 4C**). These results suggest that upregulation of the hedgehog signaling pathway within the udNSC population may be responsible for at least part of the effect of deletion of PPM1K protein.

STRING analysis of the top 30 upregulated genes within the udNSC population confirmed the hedgehog signaling node along with associated downstream genes (Ccnd1and Ccnd2), while the top 30 downregulated genes identified metallothioneins as the top hits (**Figures 4E and S4B**). To follow the perturbation of the hedgehog signaling pathway along the lineage trajectory of the four major cell types in our cultures, we generated a heatmap of the primary hedgehog pathway gene-set over the four cell types and computed average expression and fold-change difference between control and *Ppm1k* knockouts (**Figure 4F**). These heatmaps revealed clear distinctions in the lineage expression of genes that included Gli1, Ptch1,Ccnd1, and Ccnd2 in udNSCs of *Ppm1k* knockouts compared to controls.

While the highest average expression for Gli1, Ptch1, Ccnd1, and Ccnd2 occurred in the iNeu and TA populations in the WT condition, the population with the highest average expression shifted to the udNSC population in *Ppm1k* knockout cultures for Gli1, Ptch1, and Ccnd1 (**Figure 4F**). Additionally, the largest fold change consistently occurred in the udNSC population for each gene. Violin plots of Gli1 and Ptch1 reveal a higher proportion of knockout cells express Gli1, Ccnd1, and Ptch1 (**Figure 4G**). That we found the largest differences in the expression of hedgehog pathway genes occurring in the udNSC population of *Ppm1k* knockout cells is consistent with the idea that perturbation in this pathway could be causative of the activation of neural stem cells and transition to differentiating neuronal progenitors.

### Role of mTOR in the effects of PPM1K deletion on NSCs

Considering that activating BCAA metabolism through BCKDH dephosphorylation is a well described function of PPM1K, we performed experiments to determine the activity of the BCKDH enzyme in neural stem cell cultures. After 48 hours post-*Ppm1k* deletion and replenishment with fresh media, NSCs lacking PPM1K showed a pronounced reduction of BCKDH phosphorylation in control NSCs compared to those lacking PPM1K (**Figure 5A**). This result highlights the important role of PPM1K in stimulating activation of BCKDH in neural stem cells. Since levels of BCAAs are elevated in brain of *Ppm1k* cKO mice, we next wondered whether signaling pathways normally affected by BCAAs were responsible for the effects of *Ppm1k* deletion in NSCs. Activation of the mTOR pathway by leucine is a well-appreciated mechanism by which BCAAs exert some of their physiological effects (Saxton and Sabatini, 2017). Consistent with a role for elevated leucine in mTOR activation, we found rapamycin-sensitive increases in pS6K levels in *Ppm1k* KO cells compared to controls (**Figure 5B**). Importantly, activation of mTOR has been shown to crosstalk with hedgehog signaling, providing a potential mechanistic connection between BCAA catabolism and the top upregulated pathway in our GSEA analysis (Wang et al., 2012; Wu et al., 2017). Consistent with this potential connection, we found that the upregulation of *Gli1* and *Ccnd1* gene expression in response to *Ppm1k* deletion was blunted in the presence of rapamycin (**Figure 5C**). Importantly, activation of NSCs, identified by *Ascl1* expression, was similarly blunted, along with genes upregulated during NSC activation such as *Fabp7*, while levels of *Neurod1* were dramatically reduced (**Figure 5C**). Additionally, *Apoe* was decreased by ∼20% in knockout cells and was not rescued by the rapamycin treatment (**Figure 5C**). These results suggest that elevated levels of BCAAs in *Ppm1k* KOs promote activation of hedgehog signaling in NSCs at least in part via mTOR activation.

**Figure 5).**
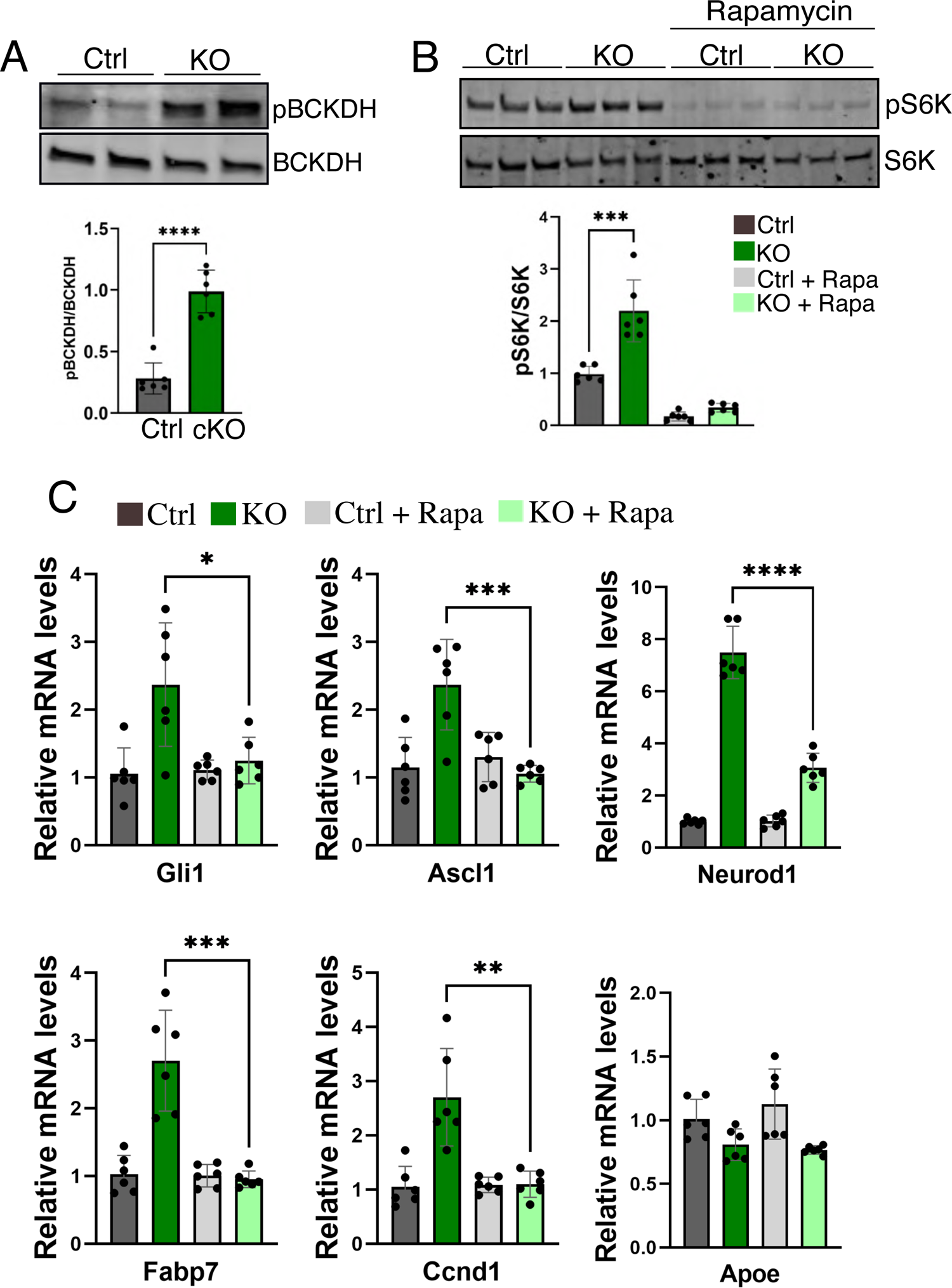
Gli1 upregulation and neural stem cell activation in *Ppm1k* KOs can be blunted by mTOR inhibition. A) Western blots for pBCKDH and BCKDH from lysates from control and *Ppm1k* KO NSC cultures in proliferation media collected 2 hours after refeeding. Quantification of the protein levels of pBCKDH presented as a fraction of total BCKDH. Ctrl n = 6, KO, n = 6; ****p < 0.0001. B) Western blots for pS6K and S6K from lysates from control and *Ppm1k* KO neural stem cell cultures in untreated of rapamycin treated condition. Quantification of the protein levels of pS6K as a fraction of S6K. Ctrl n = 6, KO, n = 6; ***p < 0.005. C) Quantification of relative mRNA expression levels for the indicated genes between control and *Ppm1k* KO NSCs either untreated or treated with 25 nM rapamycin for 36 hours. Ctrl n = 6, KO n = 6; *p < 0.05, **p < 0.01, ***p < 0.001, ****p < 0.0001. All p values were determined by independent samples two-tailed Student’s t-test

## DISCUSSION

In this study we observed a significant impairment in spatial working memory after nervous system specific deletion of the *Ppm1k* gene. We combined immunohistochemical analysis of mouse brains with single cell RNA sequencing of NSCs to find that neurogenesis was dysregulated in conjunction with incomplete maturation of newborn neurons within the dentate gyrus. Furthermore, our proliferation analysis of the dentate gyrus revealed a trend toward loss of adult neural stem cells, likely arising from their enhanced activation and differentiation. The results from these experiments combined with gene pathway analysis link the *Ppm1k* gene to working memory, hippocampal neurogenesis, and mTOR and hedgehog signaling. Our metabolomic and biochemical analyses revealed disrupted BCAA catabolism in the brains of PPM1K cKO mice and within *in vitro* neural stem cell cultures. We further identify defective BCAA catabolism as a mechanism contributing to mTOR-dependent activation of hedgehog signaling. Unlike reports that describe a reduction or increase in neurogenesis as the driver of either reduced or enhanced cognitive abilities, our observation of the high numbers of DCX+/NeuroD1 positive cells accumulating in the GCL provides a new mechanism for how defective macronutrient metabolism can lead to disruptions in working memory.

Hippocampal neurogenesis facilitates several essential hippocampal-dependent behaviors that include working memory, episodic memory, spatial memory, and mood (Clelland et al., 2009; Luna et al., 2019; Sahay et al., 2011; Toda et al., 2019). The dentate gyrus is an important region for processing spatial information, and newborn neurons are integrated into this network (Clelland et al., 2009; Kee et al., 2007; Toda et al., 2019). A recent study has shown that just a few hundred dysregulated adult-born immature neurons can cause aberrant activity in CA3 and CA1 hippocampus, leading to brain-wide disruptions in neural connectivity that impair spatial memory (Bao et al., 2020). Consistent with this finding, in this study we observed extensive accumulation of DCX+/Neurod1+ immature cells in the GCL of *Ppm1k* cKO mice that could contribute to disturbances in spatial working memory. As immature granule cells within the dentate gyrus develop, they send axons to connect to CA3 neurons while their dendrites extend into the molecular layer to receive input from the entorhinal cortex by the perforant pathway. CA3 neurons are synapse on CA1 neurons thereby completing the tri-synaptic circuit. Newborn neurons become functionally connected to the tri-synaptic circuit in approximately 4-6 weeks, though at this time they do not display electrophysiological properties similar to fully mature neurons and they contain lower thresholds and specificity during activation (Marin-Burgin et al., 2012). These immature neurons provide a level of plasticity and control to their neighboring mature neurons (Schmidt-Hieber et al., 2004). Therefore, immature neurons within the dentate gyrus provide a unique role in the tri-synaptic circuit; however it is unclear at this time how the abnormal accumulation of immature cells within the GCL can disrupt this circuit and affect memory processes. Defects in spatial working memory due to the accumulation of DCX+/Neurod1+ neurons within the dentate gyrus in *Ppm1k* cKO mice could occur at multiple points along the circuit. One possibility is that the extensive dendritic structures that are DCX-label positive in *Ppm1k* cKO mice could lack proper synaptic maturation to receive input from the entorhinal cortex, while the axons that extend through the mossy fibers may lack mature integration properties to communicate effectively with CA3 neurons. The outcome of this incomplete maturation would render thousands of PPM1K deficient neurons that reside within the granular cell layer of the dentate gyrus incapable of contributing to normal function within the tri-synaptic circuit. Future studies using behavioral and genetic approaches are required to address how spatial memory and underling circuits may be disrupted in *Ppm1k* cKO mice.

The hedgehog family of signaling molecules are involved in a wide variety of processes from developmental body plan segmentation to proliferation, differentiation, and aging (Sasai et al., 2019). Some adult tissues, such the brain’s neurogenic zones, continue to utilize hedgehog signaling for maintenance of ongoing cell fate and differentiation decisions. Sonic hedgehog signaling activates quiescent neural stem cells and transit amplifying cells in the subventricular zone and dentate gyrus, stimulating proliferation and contributing to generation of newborn cells within both olfactory bulbs or the granule cell layer of the dentate gyrus (Ahn and Joyner, 2005). Genetic methods to increase hedgehog activity, as in patch heterozygous mice, lead to increased progenitor activation and accumulation of transit amplifying cells that are Sox2 positive (Antonelli et al., 2019). Continued activation of hedgehog signaling also leads to depletion of activated neural stem cells and reduced neurogenesis in the long-term (Daynac et al., 2016). Hedgehog signaling has also been shown to regulate multiple steps in the differentiation of neurons, from subtype specification, axon guidance, synapse formation and plasticity (Belgacem et al., 2016). While hedgehog signaling induces differentiation of ventral neuronal subtypes in the developing forebrain, it also promotes differentiation of postmitotic neurons and regulates aspects of neuron identity through regulation of neuronal activity and calcium spikes (Belgacem and Borodinsky, 2011). In our studies, we observed activation of neural stem cells, depletion of stem cell number *in vivo*, and disrupted differentiation of granule cell neurons, consistent with elevated hedgehog signaling in the absence of PPM1K.

A possible activator of hedgehog signaling in *Ppm1k* cKO NSCs could be the mTOR pathway via pS6K-mediated phosphorylation of Gli1 on Ser84, resulting in its release from Sufu-mediated inhibition (Wang et al., 2012). Additional connections between mTOR and hedgehog have been reported through regulation of 4EBP1, suggesting a close association between the nutritional sensing pathway and cell fate and differentiation (Wu et al., 2017). The BCAA leucine is a well-known activator of mTOR and was increased by 40% in *Ppm1k* cKO brains, a rise that may be sufficient to activate mTOR/hedgehog signaling (Saxton and Sabatini, 2017; Wolfson et al., 2016). mTOR activity is important for the maintenance and differentiation of neural stem cells and our experiments with rapamycin confirm its contribution to the activation of *Ppm1k*-depleted neural stem cells (Meng et al., 2018). mTOR activity is also important for dendrite formation and its inhibition blocks the outgrowth of dendrites *in vitro* (Skalecka et al., 2016). Overall, our results emphasize an important mechanism by which adult neural stem cells become aberrantly activated by *Ppm1k* KO-mediated elevation of BCAA, leading to persistent activation of mTOR and hedgehog pathways.

Since Nestin-Cre is constitutively expressed beginning at E12, our mouse model would predict the consequences of lifelong disruption of BCAA catabolism, specifically in the nervous system. Individuals with reductions in BCAA activity through mutations in BCKDH or PPM1K are present in the general population and these patients present with varying levels of phenotypical severity (Oyarzabal et al., 2013; Strauss et al., 2020). While whole-body absence of *Ppm1k* in mice may contribute to cardiometabolic phenotypes, functional descriptions of PPM1K effects on neurocognitive function have not been reported (Li et al., 2017; Sun et al., 2016; Walejko et al., 2021; Wang et al., 2016). The nervous system specific *Ppm1k* knockout mouse offers an ideal model where BCAA activity is reduced, but not enough to affect viability and general physical health. Importantly, these cKO mice retained normal levels of circulating BCAA concentrations. Additionally, our *Ppm1k* cKO mice maintained normal overall motor activity. While gait stance times were slightly reduced, they were accompanied by increased swing times and equivalent stride lengths, running speeds, gait angles, distance traveled, and stride couplings.

Maple syrup urine disease (MSUD) is a developmental disorder caused by whole body reduction in catabolism of BCAAs that can be rescued by liver transplantation or dietary restriction of BCAA intake (Nellis and Danner, 2001). Adults and children with MSUD display various neuropathologies, that are often attributed to high circulating levels of BCAAs and their ketoacids, producing toxicity via accumulation of organic acids. However, despite these known neuropathologies, intrinsic nervous system-specific mechanisms for MSUD-related neurologic phenotypes are poorly defined (Xu et al., 2020). Because the circulating levels of BCAAs in our *Ppm1k* cKO mice were not altered, our mouse model allows us to remove the contribution of increased circulating levels of BCAAs and ketoacids that have been proposed to underlie the neuropathology seen in humans, and specifically uncover the contribution of reduced nervous system BCAA catabolism. BCAA levels in *Ppm1k* cKO brains remained in the low micromolar range, making it unlikely that toxicity is contributing to the impairments in working memory.

Nuclear bomb test-derived ^14^C dating has shown that up to one third of human hippocampal neurons are subject to turnover in a lifetime at a rate of 1.7% per year (Spalding et al., 2013). Neuronal cells within the dentate gyrus make up the bulk of this turnover, revealing a significant capacity for plasticity and restructuring within the neurogenic region of the hippocampus. Our results demonstrate an important role for PPM1K in regulating hippocampal neurogenesis, where impaired neuronal BCAA metabolism translates into a marked defect in spatial working memory. While dysregulated BCAA metabolism is implicated in a variety of cardiometabolic diseases (White et al., 2018), to our knowledge, this is the first study to show the contribution of neuronal BCAA metabolic perturbations to cognitive dysfunction, independent of systemic changes in BCAA metabolism. These results additionally predict that individuals genetically-predisposed to chronically insufficient BCAA catabolism may acquire deficiencies in hippocampal-dependent memory that could be alleviated by pharmacologically enhancing BCAA catabolism or by reducing mTOR activity.

## ACKNOWLEDGEMENTS

We thank the Duke University School of Medicine for the use of the Sequencing and Genomic Technologies Shared Resource, which provided single-cell RNA sequencing service, and Vaibhav Jain for assistance with data analysis. We thank the Duke Molecular Physiology Institute Metabolomics Core, which provided metabolomics service. We thank Scott Soderling and Chris Newgard for critical discussion of the project. This work was partially supported by National Institutes of Health grant K08HL135275 (to R.W.M.), American Diabetes Association Pathways to Stop Diabetes Initiator Award 1-16-INI-17 (to P.J.W.), Duke University School of Medicine Strong Start Award (to R.W.M.) and Diabetes and Endocrine Research Center grant P30 DK124723 (to Duke Molecular Physiology Institute Metabolomics Core).

## AUTHOR CONTRIBUTIONS

Conceptualization, K.A., R.W.M., P.J.W.; Methodology, K.A., R.M.R., W.C.W., R.W.M., P.J.W.; Investigation, K.A., R.M.R., W.C.W., M.E.A.; Writing – Original Draft, K.A.; Writing – Review & Editing, K.A., W.C.W., R.W.M., P.J.W; Supervision, R.W.M., P.J.W.; Funding Acquisition, R.W.M., P.J.W.

---

Figure S1. Quantification of absolute amino acid levels in brain tissue Quantification of the absolute concentration of amino acids from control and *Ppm1k* cKO brain tissue.

Figure S2. Open field activity, gait responses, and radial maze A) Locomotor activity (distance traveled in cm) in the center zone of the open field in 5-min segments over 60 min. *Inset*: cumulative center distance traveled. No significant difference. Ctrl n = 9, cKO n = 10. B) Time (sec) spent in 5 min segments in the center zone of the open field over 60 min. *Inset*: cumulative center time. No significant difference. Ctrl n = 9, cKO n = 10. C) Front and rear stride width in the gait test. Fore paw width was narrower in mutants than controls. Ctrl n = 10, cKO n = 10; *p < 0.05, multivariate ANOVA. D) Stride length with the fore paws and rear paws in the gait test. No significant difference. Ctrl n = 10, cKO n = 10. E) Front paw and rear paw gait angle in the gait test. No significant difference. Ctrl n = 10, cKO n = 10. F) Homolateral, Homologous, and Contralateral coupling in the gait test. No significant difference. Ctrl n = 10, cKO n = 10. G) Running speed in the gait test. No significant difference. Ctrl n = 10, cKO n = 10. H) Total numbers of arms used in the radial maze. Ctrl n=14, cKO n=9; p = 0.077, independent samples two-tailed Student’s t-test. I) Total numbers of perseverative errors in the radial maze. Ctrl n=14, cKO n=9; **p < 0.05, independent samples two-tailed Student’s t-test. J) Numbers of arm entries after the first error in the radial maze. Ctrl n=14, cKO n=9; *p < 0.05, independent samples two-tailed Student’s t-test. K) Entries to first repeat in the radial maze. No significant difference. Ctrl n = 14, cKO n = 9.

Figure S3. Analysis of brain anatomy and neurogenesis in PPM1K KO vs controls A) Cresyl violet stain of coronal brain sections from PPM1K^Fl/Fl^ and Nestin-Cre;PPM1K^Fl/Fl^ mice. B) Western blots showing levels of pTau and Tau in control and PPM1K cKO brains. Positive controls for pTau were acquired from brain lysates derived from the APP;eNos2(KO/KO) Alzheimer’s mouse model (Ad Tg). Quantification of the levels of pTau as a fraction of total Tau between control and PPM1K cKO brains. Ctrl, n = 6; cKO, n = 6. C) Representative images of mRNA in situ expression signal and H&E stain for PPM1K and BCKDH of mouse brain sagittal sections acquired from Allen Brain Bank Institute. White square outlines the hippocampus with region of interest zoomed in for images on right panel. Hpp = hippocampus. DG = dentate gyrus D) Representative confocal images of the dentate gyrus from mice of the genotypes PPM1K (Fl/Fl), Cre;WT, and Cre;PPM1K (Fl/Fl) labeled with NeuN, and DAPI. Scale bars = 100 μm. E) Representative wide-field confocal images of the hippocampus from mice of the genotypes PPM1K (Fl/Fl), Cre;WT, and Cre;PPM1K (Fl/Fl) labeled with DCX, NeuN, and DAPI. Scale bars = 300 μm.

Figure S4. scRNA sequence and pathway analysis A) UMAP plot showing 16 original clusters annotated by color representation. B) STRING protein analysis of the top downregulated differentially expressed genes from the udNSC group with unconnected nodes removed. Colored circles represent the top two pathways identified, red includes genes from “metallothioneins bind metals” pathway, while blue circle represent genes from “keratan sulfate degradation pathway”.

## METHODS

### Animals

*Ppm1k*^Flox/Flox^ mice on the C57Bl6 background were generated through Cyagen Corporation. *Ppm1k*^flox/flox^ mice were crossed with Nestin-Cre mice (B6.Cg-Tg(Nes-cre)1Kln/J; The Jackson Laboratory; stock, 003771) to generate Nestin-Cre;*Ppm1k*^Flox/Flox^ experimental mice. Animals were housed in groups of 3–5 animals/cage in a temperature (22°C) and humidity (45%)-controlled room with a 14 : 10 light–dark cycle (lights on at 0700 hours). Food and water were provided *ad libitum*. All experiments were conducted during the light cycle, except the home-cage emergence test that occurred in the dark. All studies were performed in accordance with NIH guidelines for the care and use of animals and with an approved animal protocol from the Duke University Institutional Animal Care and Use Committee.

### Eight-arm radial maze spontaneous alternation test

Mice were placed into the center of an 8-arm radial maze and given free access to the maze for 5 min. All tests were video-recorded and scored subsequently by an observer who was blinded to the genotypes of the mice (Kim et al., 2013). Entry into an arm was defined as the mouse being more than 1 body length into that arm, with both hind-paws past the entrance to that arm. An arm alternation was defined as successive entries into one of the different arms. Perseveration was defined as a re-entry into the previous arm. Alternation was calculated as the total number of alternations divided by the total number of arm entries minus 7 and expressed as a percentage. Variables measured include; number arms used out of the 8 available arms (Arms used), number of arm entries including repetitive entries (Arm entries), total number of re-entries (Total errors), latency time before first error (Latency 1^st^ error), ratio of the total errors over arms used (% Revisits/Arms used).

### Gait analysis - Treadscan (Clever Sys) behavior test

Analyses of gait was performed with the TreadScan apparatus and software (CleverSys Inc., Reston, VA) that generated measurements of the four phases of gait (stance, swing, propel, brake), stride length and width, paw area, toe spread, diagonal coupling (homolateral, homologous, contralateral), gait angle, body rotation, and running speed (Wang et al., 2011).

### Open field test

Spontaneous activity in the open field was conducted over 60 min in an automated Omnitech Digiscan apparatus (AccuScan Instruments, Columbus, OH) as previously described (Kim et al., 2013; Pogorelov et al., 2005). Accuscan software scored the total distance traveled, vertical activity (beam-breaks), distance traveled in the center zone, and time spent in the center zone were scored with Accuscan software in 5-min intervals.

### Novel object recognition

Mice were examined for short-, long-term, and remote memory. Testing was conducted over 5 min in four phases; object training, short-term recall at 20 min, long-term recall at 24 hr, and remote recall at 10 days. At training, mice were exposed to a pair of identical objects (2 x 2 x 3 cm in size; Iwako Inc. USA, San Francisco, CA) affixed with double-sided tape to the floor of a white Plexiglas arena (41 x 18 x 30 cm); these objects constituted the “familiar” objects for the tests. In the short- and long-term memory tests, one of the two familiar objects was replaced with a novel object with similar dimensions as the former but with different colors, patterns, and shapes. All tests were filmed with digital cameras and the videos were analyzed with Noldus Ethovision software that automatically tracked the location of each animal as well as the location and movement of the animal’s head and nose during testing. From these data, the total numbers of contacts and durations of object contacts were measured. Orientation and time spent with objects was defined as the animal’s head oriented towards the object with the nose positioned within 2 cm of the object. Recognition scores were calculated by subtracting the time spent with the familiar from the time spent with the novel object, and dividing this difference by the total time spent with both objects. Positive scores signified recognition of the novel object, negative scores indicated preferences for the familiar object, and scores approaching ‘zero’ denoted preference for neither object.

### Adenoviral constructs

Adeno-CMV-Cre-IRES-GFP and Adeno-CMV-IRES-GFP were generated and produced in 293 as previously described (Haldeman et al., 2019).

### Immunohistochemistry

Mouse brains were fixed using 4% PFA solution, equilibrated in 25% sucrose and embedded in OTC medium then frozen in −80 °C. OTC embedded mouse brains were coronally sectioned at 25 microns and mounted on Fisherbrand 24mm/74mm Superfrost Plus slides. Brain sections were permeabilized using PBS containing 0.2% triton-X100 (PBST) for one hour, blocked in PBST containing 4% BSA for 1 hour. Neural stem cell cultures were fixed in 3% PFA for 30 minutes followed by permeabilization in PBS containing 0.1% triton-X 100 in PBS for 30 minutes, then blocked in PBST containing 4% BSA for 1 hour. Primary antibodies were incubated overnight at 4 ° C and secondaries were incubated for 3 hours at 4 °C. Images were acquired on Leica TCS SP8 confocal microscope.

### Neural stem cell cultures

Primary neural stem cell cultures from p28 mice were prepared as previously described with few modifications (Wachs et al., 2003). Briefly, dentate gyrus brain tissue containing neural stem cells were mechanically minced and triturated, then digested in papain and DNase in Neurobasal A medium. A preincubation step was performed on 6 well plates to remove astrocytes, and non-adherent neural stem cells were pelleted and replated in 24 well plates. Cells were expanded in Neurobasal A containing,, B-27, glutamax, pen/strep, 10% FBS, 20 ng/ml FGF, 20 ng/ml EGF. Neural stem cell cultures were passaged using Trypsin-EDTA, expanded, and frozen for further use. To knockout PPM1K in cell cultures, adenovirus containing the pAdeno-CMV-Cre-IRES-GFP construct was delivered to subconfluent neural stem cells cultures in growth media. After 12 hours media was replaced with fresh growth media and cells were grown for at least 48 hours longer. For imaging neural stem cell cultures were grown on poly-D-lysine coated 12 mm round coverslip placed in 24 well culture dishes. For BCAA treatments neural stem cell cultures were placed in DMEM media (Genaxxon bioscience, C4150) containing 10 mM glucose, B-27 supplement, pen/strep, essential amino acids (added at concentrations normally found in DMEM) but lacking BCAAs, which were supplemented at the indicated concentrations with or without leucine. For rapamycin treatment experiments neural stem cell cultures fist treated with adenoviruses containing Cre for 16 hours in growth conditions then switch to growth media containing 25 nM rapamycin for additional 36 hours.

### Western blots

Tissue and cell lysates were prepared in Cell Lysis Buffer containing HALT protease and phosphatase-inhibitor cocktail (ThermoFisher Scientific). Samples were run on 15 well Mini-Protean TGX 4%– 15%, Stain-Free precast gels (Bio-Rad) and transferred to LF-PVDF membranes using the Trans-blot turbo system (Bio-Rad). PVDF membranes were blocked using 5% BSA in Tris-Buffered Saline buffer. Primary antibodies were incubated with membranes overnight at 4 °C and secondary antibodies were incubated for 2 hours at room temperature. Membranes were exposed and captured using a Li-Cor Odyssey CLx.

### Antibodies

Rabbit DCX (Cell Signaling #4604); mouse NeuN (abcam ab104224); rabbit NeuN (Proteintech #26975-1-AP); chicken GFAP (Novus Biologicals #NBP1-05198); rabbit NeuroD1 (Proteintech #12081-1-AP); rabbit Sox2 (Genetex #GTX101507); rabbit BCKDH, (abcam #ab126173); rabbit pBCKDH (abcam #ab200577); rabbit PPM1K (Proteintech #14573-1-AP); rabbit actin (abcam #ab1801); mouse gTub, (Sigma T6557); rabbit pS6K (Cell Signaling #9234); rabbit S6K, (Cell Signaling #2708). Mitotracker was purchased from Thermo Fischer (M7512).

### Amino acid, acylcarnitine, ketoacids measurements

Targeted amino acid, acylcarnitine, and branched-chain α-ketoacid analyses were performed from brain tissue with internal standards as previously described (Newgard, 2012; White et al., 2018).

### Single cell RNA sequencing

*Ppm1k*^Flox/Flox^ and *Ppm1k*^Wt/Wt^ neural stem cell cultures were treated with CMV-Adeno-Cre-IRES-GFP for 68 hours in growth media then collected through GFP FACs sorting. Samples were prepared using the General Sample Preparation protocol from 10× Genomics (10×, Manual Part #CG00053) adapted from published methods (Zheng et al., 2017). 10x Transcriptome library prep: Briefly, single cells are dissociated, then washed and resuspended in a 1x PBS / 0.04% BSA solution, at a concentration of 1000 cells/ul. After size selection (<50um), the cell suspension is washed with a 1x PBS / 0.04% BSA solution to remove debris, clumps, dead cells and contaminants, and a Cellometer (Nexcelom - Lawrence, MA) is used to determine the cell viability and concentration to normalize to 1×10^6^ cells/ml. We then titer each prep to contain ∼10,000 cells per library construction. cDNAs are assayed on an Agilent 4200 TapeStation High Sensitivity D5000 ScreenTape (Santa Clara, CA) for qualitative and quantitative analysis. Enzymatic fragmentation and size selection are used to optimize the cDNA amplicon size. Illumina P5 and P7 sequences (San Diego), a sample index, and TruSeq read 2 primer sequence are added via End Repair, A-tailing, Adaptor Ligation, and PCR. The final libraries contain P5 and P7 primers used in Illumina bridge amplification. Sequence is generated using paired end sequencing (one end to generate cell specific, barcoded sequence and the other to generate sequence of the expressed poly-A tailed mRNA) on an Illumina sequencing platform at a minimum of 50k reads/cell.

*Analysis*: The primary analytical pipeline for the SC analysis follows the recommended protocols from 10X Genomics. Briefly, we demultiplex raw base call (BCL) files generated by Illumina sequencers into FASTQ files, upon which alignment to the appropriate reference transcriptome, filtering, barcode counting, and UMI counting are performed using the most current version of 10X’s Cell Ranger software. The secondary statistical analysis is performed using the R package Seurat, which performs quality control and subsequent analyses on the feature-barcode matrices produced by Cell Ranger. In Seurat, data is first normalized and scaled after basic filtering for minimum gene and cell observance frequency cut-offs. We then closely examine the data and perform further filtering based a range of metrics in attempt to identify and exclude possible multiplets (i.e. instances where more than one cell was present and sequenced in a single emulsified gel bead). The removal of further technical artifacts is performed using regression methods to reduce noise. We perform linear dimensional reduction calculating principal components using the most variably expressed genes in our dataset. Library size and/or the numbers of genes expressed across subsets of cells may necessitate the restriction of cells upon which the variably-expressed genes are selected for inclusion when calculating principal components. The genes underlying the resulting principal components are examined in order to confirm they are not enriched in genes involved in cell division or other standard cellular processes. Significant principal components for downstream analyses are determined through methods mirroring those implemented by Macosko et al. Cells are grouped into an optimal number of clusters for de novo cell type discovery using Seurat’s FindNeighbors() and FindClusters() functions, graph-based clustering approach with visualization of cells being achieved through the use of t-SNE, which reduces the information captured in the selected significant principal components to two dimensions. Differential expression genes were identified using a Wilcoxon Rank-Sum test with a cut-off of +/- 0.25 log2fold change and adj-P <0.05.

### qPCR

Total RNA was prepared from mouse neural stem cell cultures and mouse brain tissue using Qiagen RNeasy kit. RNA was reverse transcribed using Multiscribe from Invitrogen (Cat#-4311235). qPCRs were run using PowerUp SYBR green master mix on the QuantStudio 6 Flex Real-Time PCR system (Applied Biosystems). Mouse primers used for qPCR:

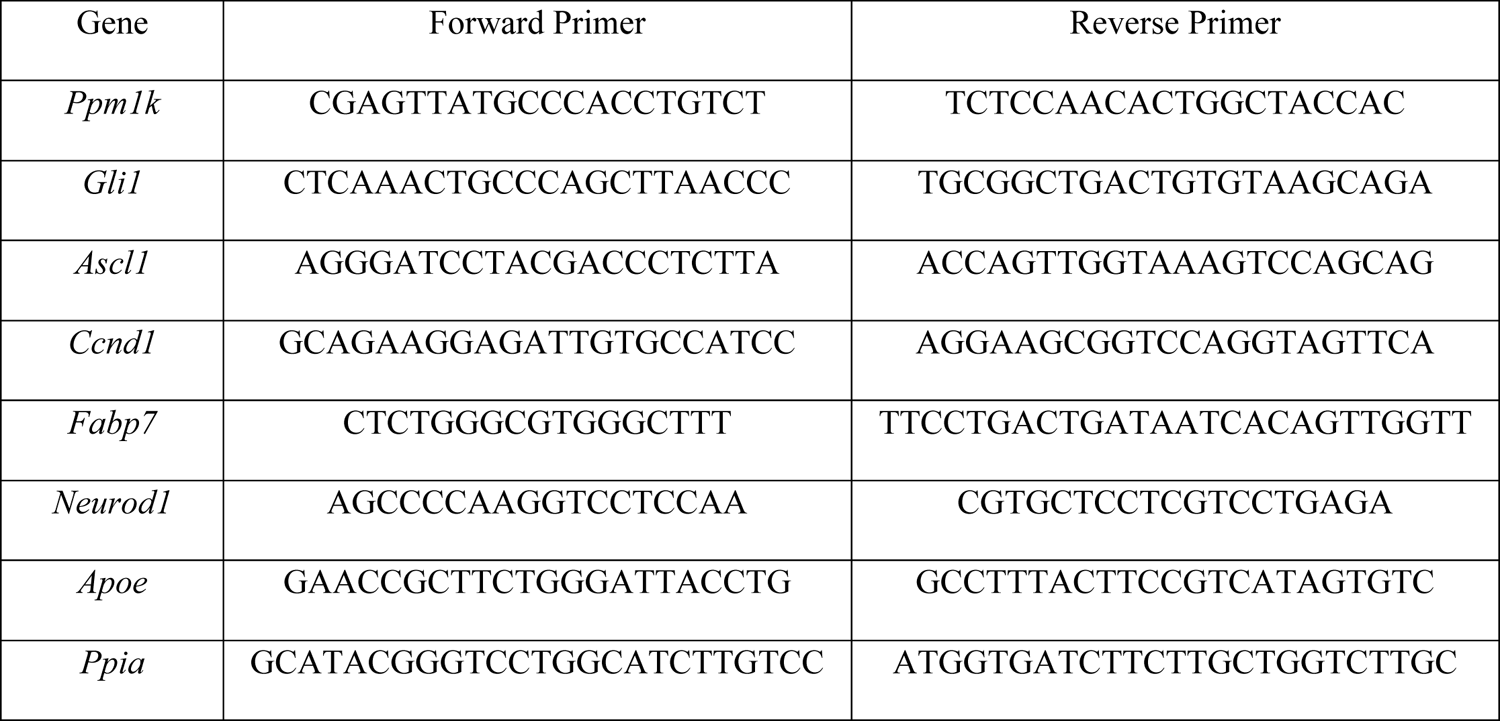

### Pathway analysis

Gene set enrichment analysis was performed using the GSEA_4.1.0 software downloaded from the Broad Institute (https://www.gsea-msigdb.org/gsea/index.jsp). An FDR of 0.1 and p values < 0.05 was used as a cutoff for generating lists of upregulated or downregulated pathways. Over-representation analysis was performed using software from Webgestalt using a pre-ranked list of differentially expressed genes with p values < 0.01 (http://www.webgestalt.org/). Heatmaps were generated using list of genes identified in GSEA pathways and their normalized expression values in each cell type with the Morpheus software from the Broad Institute (https://software.broadinstitute.org/morpheus/). STRING analysis was performed using the STRING online software found through the Elixir Core Data resource site (https://string-db.org/).

## Statistical Analysis

Figures were presented as mean ± standard deviation (SD). For representative images, western blots, and associated quantifications the results were shown to be reproducible by at least three separate experiments. Significance between two conditions was analyzed using Student’s t-test and individual data points are presented in graphs to show variations from mean. The behavioral data were analyzed with SPSS statistics (IBM) and presented as means and standard error of the mean. Group differences were analyzed by t-tests, repeated measures ANOVA (RMANOVA), or multivariate ANOVA (MANOVA) followed by Bonferroni corrected pairwise comparisons. These statistics for the different behavioral tests are presented in Supplementary Table S1 & S2. A p < 0.05 was considered statistically significant. For differential expression a Wilcoxon rank sum test was performed and included cutoffs for log2 fold changes of +/- 0.25 and p < 0.05.

