## Supplementary Material for "Nervous system reduction in branched-chain amino acid metabolism disrupts hippocampal neurogenesis and memory"

### Figure S1

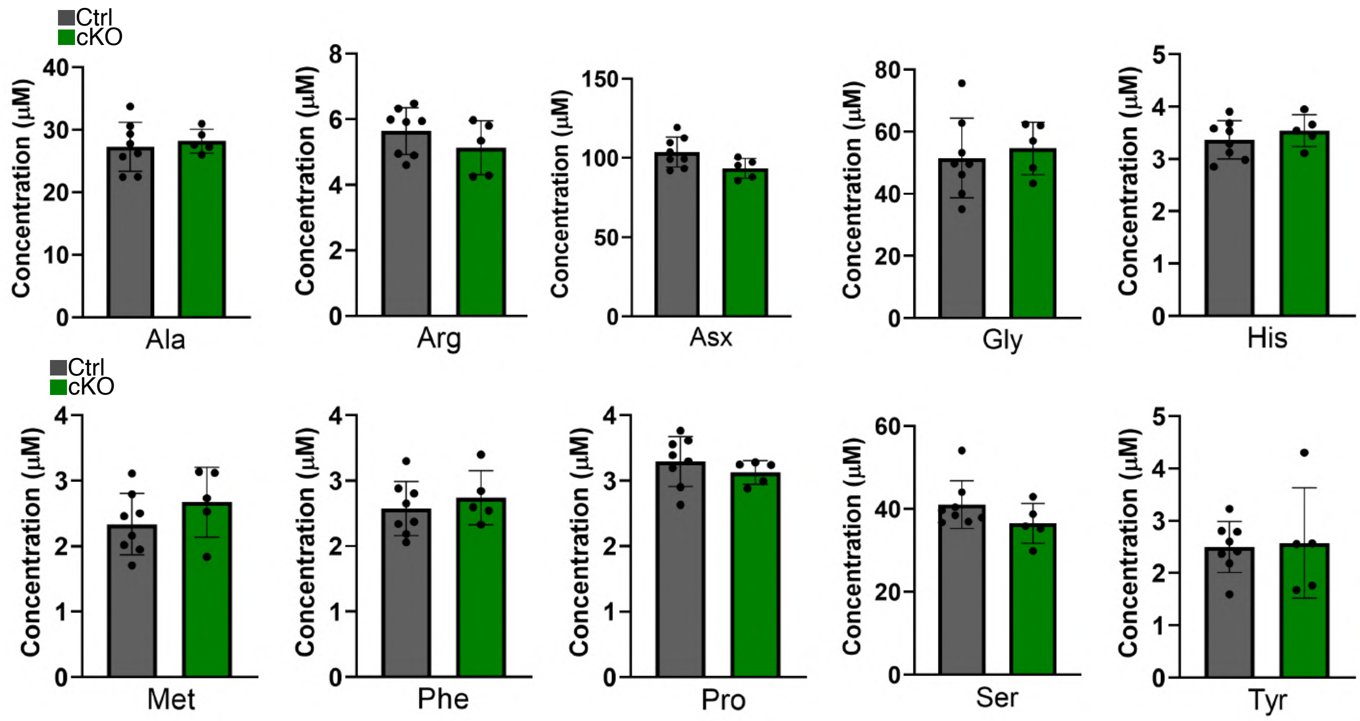
Ctrl
cKO
Concentration (μM)
Ala
Arg
Asx
Gly
His
Met
Phe
Pro
Ser
Tyr

### Figure S2

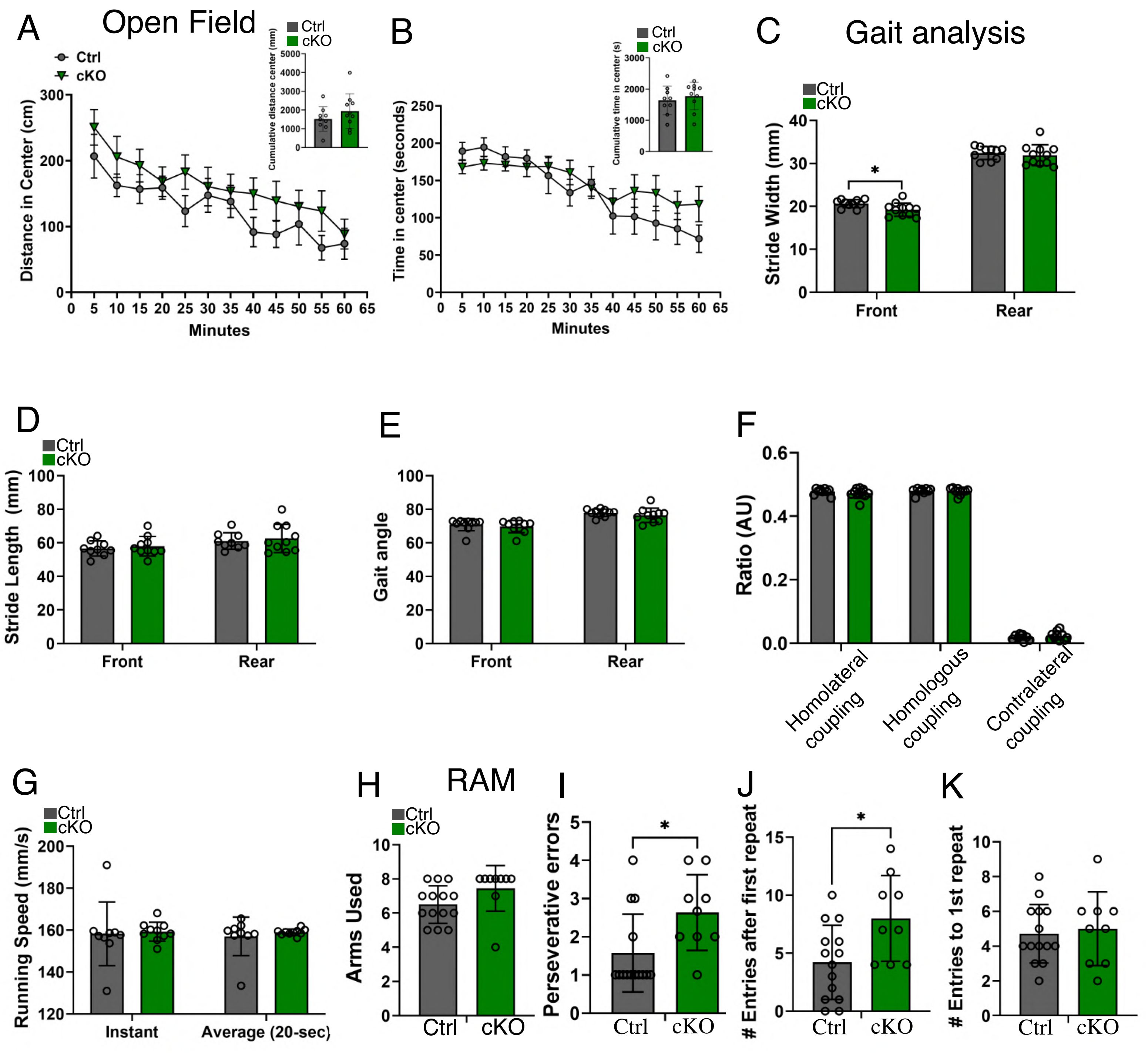

A
Open Field
Ctrl
cKO
Distance in Center (cm)
Minutes
Cumulative distance center (mm)
B
Time in center (seconds)
Cumulative time in center (s)
C
Gait analysis
Stride Width (mm)
Front
Rear
D
Stride Length (mm)
E
Gait angle
F
Ratio (AU)
Homolateral coupling
Homologous coupling
Contralateral coupling
G
Running Speed (mm/s)
Instant
Average (20-sec)
H
RAM
Arms Used
I
Perseverative errors
J
### Entries after first repeat
K
### Entries to 1st repeat

### Figure S3

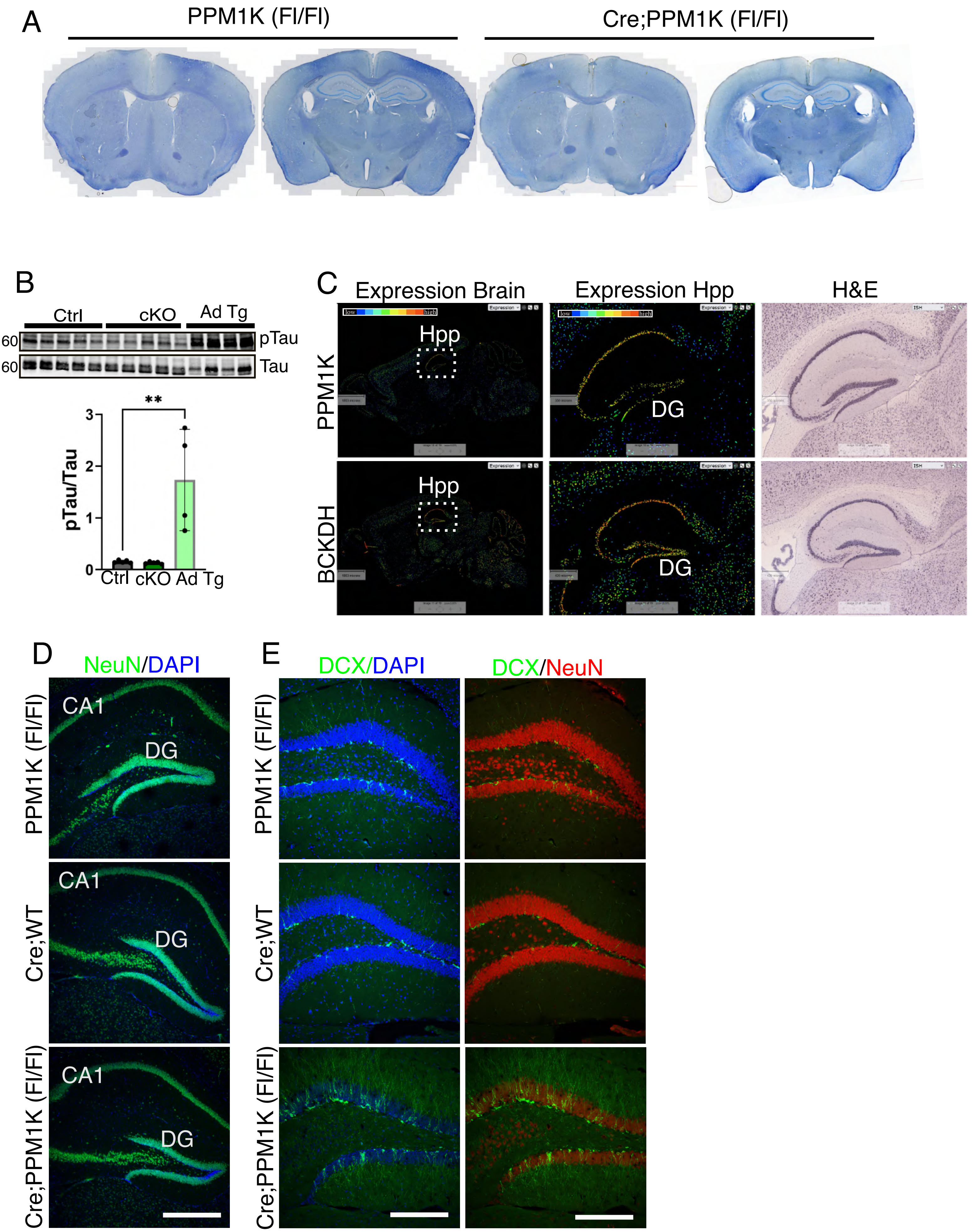
A
PPM1K (Fl/Fl)
Cre;PPM1K (Fl/Fl)
B
Ctrl
cKO
Ad Tg
60
pTau
60
Tau
**
pTau/Tau
3
2
1
0
Ctrl
cKO
Ad Tg
C
Expression Brain
Expression Hpp
H&E
PPM1K
BCKDH
Hpp
DG
low
high
D
NeuN/DAPI
PPM1K (Fl/Fl)
Cre;WT
Cre;PPM1K (Fl/Fl)
CA1
DG
E
DCX/DAPI
DCX/NeuN

**Figure S4**

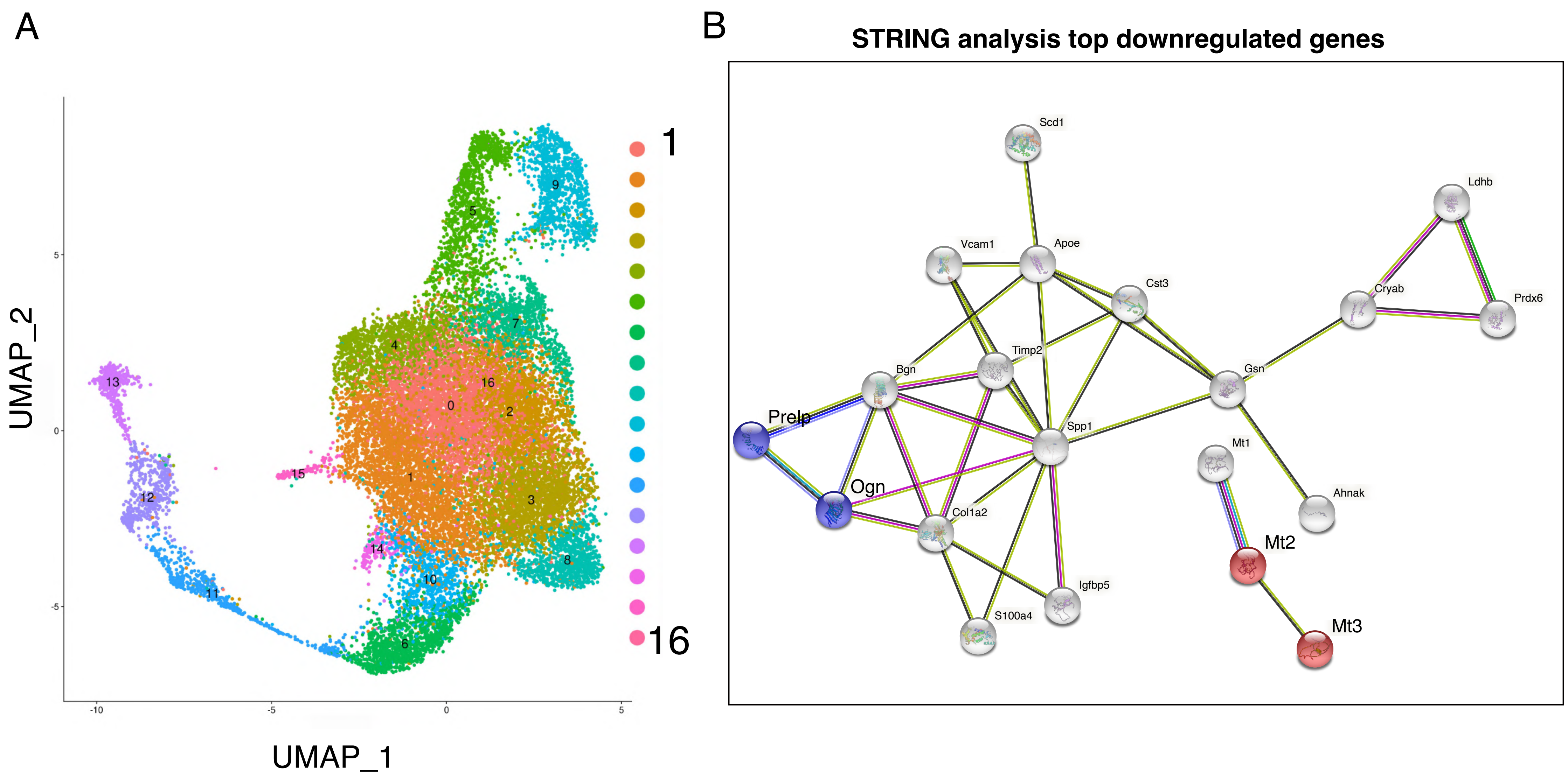
A
B
STRING analysis top downregulated genes
UMAP_2
UMAP_1
1
16
Scd1
Ldhb
Vcam1
Apoe
Cst3
Cryab
Prdx6
Timp2
Bgn
Gsn
Prelp
Spp1
Mt1
Ogn
Ahnak
Col1a2
Mt2
Igfbp5
S100a4
Mt3

**Supplementary Table S1.**

**Statistics for the Behavioral Studies in Figures**

| **Figure** | **Stat. Model** | **Variable** | **Deg. of Free.** | **F-statistic** | ***p*-value** | **#mice/gr.** |
|---|---|---|---|---|---|---|
| Fig. 1G | RMANOVA[a] | Time | 11,187 | 13.811 | <0.001 | 9 Ctrl, 10 cKO |
| | t-test | Genotype | 17 | 1.444 | n.s.[a] | 9 Ctrl, 10 cKO |
| Fig. 1H | RMANOVA | Time | 11,187 | 7.215 | <0.001 | 9 Ctrl, 10 cKO |
| | | Time x Genotype | 11,187 | 2,358 | 0.010 | 9 Ctrl, 10 cKO |
| | t-test | Genotype | 17 | 1.798 | 0.045 | 9 Ctrl, 10 cKO |
| Suppl. Fig. S2A | RMANOVA | Time | 11,187 | 17.505 | <0.001 | 9 Ctrl, 10 cKO |
| | t-test | Genotype | 17 | 1.151 | n.s.[a] | 9 Ctrl, 10 cKO |
| Suppl. Fig. S2B | RMANOVA | Time | 11,187 | 11.903 | <0.001 | 9 Ctrl, 10 cKO |
| | t-test | Genotype | 17 | 0.668 | n.s.[a] | 9 Ctrl, 10 cKO |
| Fig. 1I | RMANOVA | Gait-phase | 3,54 | 99.204 | <0.001 | 10 Ctrl, 10 cKO |
| | | Gait-phase x Gene | 3,54 | 7.461 | 0.007 | 10 Ctrl, 10 cKO |
| Fig. 1J | RMANOVA | Gait-phase | 3,54 | 55.814 | <0.001 | 10 Ctrl, 10 cKO |
| | | Gait-phase x Gene | 1,18 | 4.510 | 0.048 | 10 Ctrl, 10 cKO |
| Fig. 1K | MANOVA | Fore-paws | 1,18 | 4.315 | 0.052 | 10 Ctrl, 10 cKO |
| | | Hind-paws | 1,18 | 25.694 | <0.001 | 10 Ctrl, 10 cKO |
| Fig. 1L | MANOVA | Toe-spread | 1,18 | 4.059 | n.s.[a] | 10 Ctrl, 10 cKO |
| | | Inter-toe Dist | 1,18 | 14.983 | 0.001 | 10 Ctrl, 10 cKO |
| Suppl. Fig. S2C | MANOVA | Fore-limbs | 1,18 | 5.979 | 0.025 | 10 Ctrl, 10 cKO |
| | | Hind-limbs | 1,18 | 0.168 | n.s.[a] | 10 Ctrl, 10 cKO |
| Fig. 1M | t-test | Lat to 1st Error | 21 | 2.732 | 0.012 | 14 Ctrl, 9 cKO |
| Fig. 1N | t-test | # Entries | 21 | 3.578 | 0.0018 | 14 Ctrl, 9 cKO |
| Fig. 1O | t-test | # Arms Revisit | 21 | 4.110 | <0.001 | 14 Ctrl, 9 cKO |
| Fig. 1P | t-test | #Total Errors | 21 | 3.466 | 0.002 | 14 Ctrl, 9 cKO |
| Fig. 1Q | t-test | % Error/Entries | 21 | 3.607 | 0.0017 | 14 Ctrl, 9 cKO |
| Suppl. Fig. S2H | t-test | # Arms Used | 21 | 1.858 | 0.077 | 14 Ctrl, 9 cKO |
| Suppl. Fig. S2I | t-test | # Perseverative Errors | 21 | 2.460 | 0.022 | 14 Ctrl, 9 cKO |
| Suppl. Fig. S2J | t-test | # Entries After 1st Error | 21 | 2.608 | 0.016 | 14 Ctrl, 9 cKO |
| Suppl. Fig. S2K | t-test | Entries to Repeat | 21 | 0.359 | n.s.[a] | 14 Ctrl, 9 cKO |

[a]Abbreviations: RMANOVA, repeated measures ANOVA; MANOVA, multivariate ANOVA; #, number; n.s., not significant.

**Supplementary Table S2.**

**Statistics for non-significant behavioral results in gait analysis**

| Figure | Stat. Model | Variable | *p*-value | #mice/gr. |
|---|---|---|---|---|
| Fig. S2D | MANOVA | Front | n.s. | 10 Ctrl, 10 cKO |
| | | Rear | n.s. | 10 Ctrl, 10 cKO |
| Fig. S2E | MANOVA | Front | n.s. | 10 Ctrl, 10 cKO |
| | | Rear | n.s. | 10 Ctrl, 10 cKO |
| Fig. S2F | MANOVA | Homolateral | n.s. | 10 Ctrl, 10 cKO |
| | | Homologous | n.s. | 10 Ctrl, 10 cKO |
| | | Contralateral | n.s. | 10 Ctrl, 10 cKO |
| Fig. S2G | MANOVA | Instant | n.s. | 10 Ctrl, 10 cKO |
| | | Average | n.s. | 10 Ctrl, 10 cKO |

[a]Abbreviations: RMANOVA, repeated measures ANOVA; MANOVA, multivariate ANOVA; #, number; n.s., not significant.

**Supplementary Table S3. Gene list for GSEA top hits**

| Basal Cell Carcinoma | SYMBOL | TITLE | RANK METRIC SCORE | RUNNING ES | CORE ENRICHMENT |
|---|---|---|---|---|---|
| 37 | BMP2 | bone morphogenetic protein 2 [Source:HGNC Symbol;Acc:HGNC:1069] | -1.102 | -0.5752 | Yes |
| 38 | TCF7 | transcription factor 7 [Source:HGNC Symbol;Acc:HGNC:11639] | -1.266 | -0.5698 | Yes |
| 39 | WNT2 | Wnt family member 2 [Source:HGNC Symbol;Acc:HGNC:12780] | -1.325 | -0.5556 | Yes |
| 40 | AXIN2 | axin 2 [Source:HGNC Symbol;Acc:HGNC:904] | -2.061 | -0.5636 | Yes |
| 41 | PTCH1 | patched 1 [Source:HGNC Symbol;Acc:HGNC:9585] | -2.409 | -0.545 | Yes |
| 42 | WNT7B | Wnt family member 7B [Source:HGNC Symbol;Acc:HGNC:12787] | -2.517 | -0.5153 | Yes |
| 43 | WNT7A | Wnt family member 7A [Source:HGNC Symbol;Acc:HGNC:12786] | -3.075 | -0.4891 | Yes |
| 44 | GLI1 | GLI family zinc finger 1 [Source:HGNC Symbol;Acc:HGNC:4317] | -3.129 | -0.4487 | Yes |
| 45 | WNT10B | Wnt family member 10B [Source:HGNC Symbol;Acc:HGNC:12775] | -3.381 | -0.4084 | Yes |
| 46 | PTCH2 | patched 2 [Source:HGNC Symbol;Acc:HGNC:9586] | -3.4 | -0.3634 | Yes |
| 47 | WNT16 | Wnt family member 16 [Source:HGNC Symbol;Acc:HGNC:16267] | -3.4 | -0.3181 | Yes |
| 48 | LEF1 | lymphoid enhancer binding factor 1 [Source:HGNC Symbol;Acc:HGNC:6551] | -3.438 | -0.2732 | Yes |
| 49 | WNT10A | Wnt family member 10A [Source:HGNC Symbol;Acc:HGNC:13829] | -3.894 | -0.2278 | Yes |
| 50 | SHH | sonic hedgehog signaling molecule [Source:HGNC Symbol;Acc:HGNC:10848] | -4.099 | -0.1751 | Yes |
| 51 | WNT6 | Wnt family member 6 [Source:HGNC Symbol;Acc:HGNC:12785] | -4.422 | -0.1195 | Yes |
| 52 | HHIP | hedgehog interacting protein [Source:HGNC Symbol;Acc:HGNC:14866] | -4.454 | -0.0604 | Yes |

| | | | | | |
|---|---|---|---|---|---|
| 53 | FZD10 | frizzled class receptor 10 [Source:HGNC Symbol;Acc:HGNC:4039] | -4.677 | 0.0005 | Yes |

| Steroid hormone biosynthesis | SYMBOL | TITLE | RANK METRIC SCORE | RUNNING ES | CORE ENRICHMENT |
|---|---|---|---|---|---|
| 1 | UGT1A4 | UDP glucuronosyltransferase family 1 member A4 [Source:HGNC Symbol;Acc:HGNC:12536] | 3.083 | 0.1141 | Yes |
| 2 | UGT1A6 | UDP glucuronosyltransferase family 1 member A6 [Source:HGNC Symbol;Acc:HGNC:12538] | 2.674 | 0.2163 | Yes |
| 3 | UGT1A3 | UDP glucuronosyltransferase family 1 member A3 [Source:HGNC Symbol;Acc:HGNC:12535] | 2.61 | 0.3215 | Yes |
| 4 | CYP21A2 | cytochrome P450 family 21 subfamily A member 2 [Source:HGNC Symbol;Acc:HGNC:2600] | 2.265 | 0.405 | Yes |
| 5 | CYP11A1 | cytochrome P450 family 11 subfamily A member 1 [Source:HGNC Symbol;Acc:HGNC:2590] | 2.188 | 0.4914 | Yes |
| 6 | CYP1B1 | cytochrome P450 family 1 subfamily B member 1 [Source:HGNC Symbol;Acc:HGNC:2597] | 2.164 | 0.579 | Yes |
| 7 | UGT1A10 | UDP glucuronosyltransferase family 1 member A10 [Source:HGNC Symbol;Acc:HGNC:12531] | 1.855 | 0.643 | Yes |

| Neuroactive ligand receptor interaction | SYMBOL | TITLE | RANK METRIC SCORE | RUNNING ES | CORE ENRICHMENT |
|---|---|---|---|---|---|
| 121 | GALR2 | galanin receptor 2 [Source:HGNC Symbol;Acc:HGNC:4133] | -0.886 | -0.4291 | Yes |
| 122 | P2RY6 | pyrimidinergic receptor P2Y6 [Source:HGNC Symbol;Acc:HGNC:8543] | -0.904 | -0.4274 | Yes |
| 123 | HRH1 | histamine receptor H1 [Source:HGNC Symbol;Acc:HGNC:5182] | -0.912 | -0.4244 | Yes |
| 124 | EDNRA | endothelin receptor type A [Source:HGNC Symbol;Acc:HGNC:3179] | -0.941 | -0.423 | Yes |
| 125 | GRID1 | glutamate ionotropic receptor delta type subunit 1 [Source:HGNC Symbol;Acc:HGNC:4575] | -0.973 | -0.4214 | Yes |
| 126 | ADRB2 | adrenoceptor beta 2 [Source:HGNC Symbol;Acc:HGNC:286] | -0.99 | -0.4186 | Yes |
| 127 | GPR156 | G protein-coupled receptor 156 [Source:HGNC Symbol;Acc:HGNC:20844] | -1.011 | -0.416 | Yes |
| 128 | NPY1R | neuropeptide Y receptor Y1 [Source:HGNC Symbol;Acc:HGNC:7956] | -1.097 | -0.4184 | Yes |
| 129 | PTH1R | parathyroid hormone 1 receptor [Source:HGNC Symbol;Acc:HGNC:9608] | -1.168 | -0.4186 | Yes |
| 130 | GRPR | gastrin releasing peptide receptor [Source:HGNC Symbol;Acc:HGNC:4609] | -1.187 | -0.4151 | Yes |
| 131 | NPFFR1 | neuropeptide FF receptor 1 [Source:HGNC Symbol;Acc:HGNC:17425] | -1.259 | -0.415 | Yes |
| 132 | MCHR1 | melanin concentrating hormone receptor 1 [Source:HGNC Symbol;Acc:HGNC:4479] | -1.276 | -0.411 | Yes |
| 133 | MC3R | melanocortin 3 receptor [Source:HGNC Symbol;Acc:HGNC:6931] | -1.318 | -0.4077 | Yes |
| 134 | CHRNB1 | cholinergic receptor nicotinic beta 1 subunit [Source:HGNC Symbol;Acc:HGNC:1961] | -1.327 | -0.4029 | Yes |

| | | | | | |
|---|---|---|---|---|---|
| 135 | LPAR6 | lysophosphatidic acid receptor 6 [Source:HGNC Symbol;Acc:HGNC:15520] | -1.331 | -0.3978 | Yes |
| 136 | HTR2A | 5-hydroxytryptamine receptor 2A [Source:HGNC Symbol;Acc:HGNC:5293] | -1.394 | -0.3957 | Yes |
| 137 | P2RX2 | purinergic receptor P2X 2 [Source:HGNC Symbol;Acc:HGNC:15459] | -1.397 | -0.3902 | Yes |
| 138 | GABRB3 | gamma-aminobutyric acid type A receptor subunit beta3 [Source:HGNC Symbol;Acc:HGNC:4083] | -1.51 | -0.391 | Yes |
| 139 | ADRA2C | adrenoceptor alpha 2C [Source:HGNC Symbol;Acc:HGNC:283] | -1.627 | -0.3898 | Yes |
| 140 | ADRB3 | adrenoceptor beta 3 [Source:HGNC Symbol;Acc:HGNC:288] | -1.673 | -0.3848 | Yes |
| 141 | GALR3 | galanin receptor 3 [Source:HGNC Symbol;Acc:HGNC:4134] | -1.728 | -0.38 | Yes |
| 142 | PTGER2 | prostaglandin E receptor 2 [Source:HGNC Symbol;Acc:HGNC:9594] | -1.779 | -0.3748 | Yes |
| 143 | ADCYAP1R1 | ADCYAP receptor type I [Source:HGNC Symbol;Acc:HGNC:242] | -1.822 | -0.3694 | Yes |
| 144 | NTSR2 | neurotensin receptor 2 [Source:HGNC Symbol;Acc:HGNC:8040] | -1.944 | -0.3666 | Yes |
| 145 | PTGFR | prostaglandin F receptor [Source:HGNC Symbol;Acc:HGNC:9600] | -1.987 | -0.3604 | Yes |
| 146 | CYSLTR1 | cysteinyl leukotriene receptor 1 [Source:HGNC Symbol;Acc:HGNC:17451] | -2 | -0.3528 | Yes |
| 147 | MC1R | melanocortin 1 receptor [Source:HGNC Symbol;Acc:HGNC:6929] | -2.065 | -0.347 | Yes |
| 148 | GRIN2D | glutamate ionotropic receptor NMDA type subunit 2D [Source:HGNC Symbol;Acc:HGNC:4588] | -2.266 | -0.346 | Yes |
| 149 | GRIA2 | glutamate ionotropic receptor AMPA type subunit 2 [Source:HGNC Symbol;Acc:HGNC:4572] | -2.428 | -0.3416 | Yes |
| 150 | GRIN2B | glutamate ionotropic receptor NMDA type | -2.429 | -0.3316 | Yes |

| | | | | | |
|---|---|---|---|---|---|
| | | subunit 2B [Source:HGNC Symbol;Acc:HGNC:4586] | | | |
| 151 | F2RL1 | F2R like trypsin receptor 1 [Source:HGNC Symbol;Acc:HGNC:3538] | -2.485 | -0.3235 | Yes |
| 152 | SCTR | secretin receptor [Source:HGNC Symbol;Acc:HGNC:10608] | -2.486 | -0.3134 | Yes |
| 153 | GABRG3 | gamma-aminobutyric acid type A receptor subunit gamma3 [Source:HGNC Symbol;Acc:HGNC:4088] | -2.492 | -0.3033 | Yes |
| 154 | P2RY14 | purinergic receptor P2Y14 [Source:HGNC Symbol;Acc:HGNC:16442] | -2.583 | -0.2955 | Yes |
| 155 | ADORA2A | adenosine A2a receptor [Source:HGNC Symbol;Acc:HGNC:263] | -2.77 | -0.2894 | Yes |
| 156 | GABBR2 | gamma-aminobutyric acid type B receptor subunit 2 [Source:HGNC Symbol;Acc:HGNC:4507] | -2.894 | -0.2808 | Yes |
| 157 | PRSS3 | serine protease 3 [Source:HGNC Symbol;Acc:HGNC:9486] | -3.044 | -0.2717 | Yes |
| 158 | GIPR | gastric inhibitory polypeptide receptor [Source:HGNC Symbol;Acc:HGNC:4271] | -3.144 | -0.2615 | Yes |
| 159 | LTB4R2 | leukotriene B4 receptor 2 [Source:HGNC Symbol;Acc:HGNC:19260] | -3.192 | -0.2494 | Yes |
| 160 | CHRNA1 | cholinergic receptor nicotinic alpha 1 subunit [Source:HGNC Symbol;Acc:HGNC:1955] | -3.226 | -0.2369 | Yes |
| 161 | LEPR | leptin receptor [Source:HGNC Symbol;Acc:HGNC:6554] | -3.249 | -0.2238 | Yes |
| 162 | HRH2 | histamine receptor H2 [Source:HGNC Symbol;Acc:HGNC:5183] | -3.281 | -0.2108 | Yes |
| 163 | PTGDR | prostaglandin D2 receptor [Source:HGNC Symbol;Acc:HGNC:9591] | -3.298 | -0.1976 | Yes |
| 164 | GRIA1 | glutamate ionotropic receptor AMPA type subunit 1 [Source:HGNC Symbol;Acc:HGNC:4571] | -3.31 | -0.1842 | Yes |

| | | | | | |
|---|---|---|---|---|---|
| 165 | GABRE | gamma-aminobutyric acid type A receptor subunit epsilon [Source:HGNC Symbol;Acc:HGNC:4085] | -3.373 | -0.1715 | Yes |
| 166 | GRM3 | glutamate metabotropic receptor 3 [Source:HGNC Symbol;Acc:HGNC:4595] | -3.699 | -0.162 | Yes |
| 167 | HTR1B | 5-hydroxytryptamine receptor 1B [Source:HGNC Symbol;Acc:HGNC:5287] | -3.703 | -0.1469 | Yes |
| 168 | GRM5 | glutamate metabotropic receptor 5 [Source:HGNC Symbol;Acc:HGNC:4597] | -3.911 | -0.1335 | Yes |
| 169 | FPR2 | formyl peptide receptor 2 [Source:HGNC Symbol;Acc:HGNC:3827] | -3.945 | -0.1176 | Yes |
| 170 | GABRA4 | gamma-aminobutyric acid type A receptor subunit alpha4 [Source:HGNC Symbol;Acc:HGNC:4078] | -4.084 | -0.102 | Yes |
| 171 | GABRB1 | gamma-aminobutyric acid type A receptor subunit beta1 [Source:HGNC Symbol;Acc:HGNC:4081] | -4.257 | -0.0861 | Yes |
| 172 | CALCRL | calcitonin receptor like receptor [Source:HGNC Symbol;Acc:HGNC:16709] | -4.295 | -0.069 | Yes |
| 173 | CHRNA10 | cholinergic receptor nicotinic alpha 10 subunit [Source:HGNC Symbol;Acc:HGNC:13800] | -4.367 | -0.0518 | Yes |
| 174 | NPFFR2 | neuropeptide FF receptor 2 [Source:HGNC Symbol;Acc:HGNC:4525] | -4.368 | -0.034 | Yes |
| 175 | DRD2 | dopamine receptor D2 [Source:HGNC Symbol;Acc:HGNC:3023] | -4.458 | -0.0167 | Yes |
| 176 | BDKRB2 | bradykinin receptor B2 [Source:HGNC Symbol;Acc:HGNC:1030] | -4.509 | 0.0015 | Yes |

| **Hedgehog signaling** | **SYMBOL** | **TITLE** | **RANK METRIC SCORE** | **RUNNING ES** | **CORE ENRICHMENT** |
|---|---|---|---|---|---|
| 33 | WNT4 | Wnt family member 4 [Source:HGNC Symbol;Acc:HGNC:12783] | -0.904 | -0.6343 | Yes |

| | | | | | |
|---|---|---|---|---|---|
| 34 | DHH | desert hedgehog signaling molecule [Source:HGNC Symbol;Acc:HGNC:2865] | -0.916 | -0.621 | Yes |
| 35 | BMP2 | bone morphogenetic protein 2 [Source:HGNC Symbol;Acc:HGNC:1069] | -1.102 | -0.6185 | Yes |
| 36 | BMP6 | bone morphogenetic protein 6 [Source:HGNC Symbol;Acc:HGNC:1073] | -1.105 | -0.6014 | Yes |
| 37 | WNT2 | Wnt family member 2 [Source:HGNC Symbol;Acc:HGNC:12780] | -1.325 | -0.5954 | Yes |
| 38 | ZIC2 | Zic family member 2 [Source:HGNC Symbol;Acc:HGNC:12873] | -1.896 | -0.5937 | Yes |
| 39 | PTCH1 | patched 1 [Source:HGNC Symbol;Acc:HGNC:9585] | -2.409 | -0.5772 | Yes |
| 40 | WNT7B | Wnt family member 7B [Source:HGNC Symbol;Acc:HGNC:12787] | -2.517 | -0.5417 | Yes |
| 41 | WNT7A | Wnt family member 7A [Source:HGNC Symbol;Acc:HGNC:12786] | -3.075 | -0.5084 | Yes |
| 42 | GLI1 | GLI family zinc finger 1 [Source:HGNC Symbol;Acc:HGNC:4317] | -3.129 | -0.4608 | Yes |
| 43 | WNT10B | Wnt family member 10B [Source:HGNC Symbol;Acc:HGNC:12775] | -3.381 | -0.4128 | Yes |
| 44 | PTCH2 | patched 2 [Source:HGNC Symbol;Acc:HGNC:9586] | -3.4 | -0.3599 | Yes |
| 45 | WNT16 | Wnt family member 16 [Source:HGNC Symbol;Acc:HGNC:16267] | -3.4 | -0.3068 | Yes |
| 46 | BMP7 | bone morphogenetic protein 7 [Source:HGNC Symbol;Acc:HGNC:1074] | -3.72 | -0.254 | Yes |
| 47 | WNT10A | Wnt family member 10A [Source:HGNC Symbol;Acc:HGNC:13829] | -3.894 | -0.1953 | Yes |
| 48 | SHH | sonic hedgehog signaling molecule [Source:HGNC Symbol;Acc:HGNC:10848] | -4.099 | -0.1332 | Yes |
| 49 | WNT6 | Wnt family member 6 [Source:HGNC Symbol;Acc:HGNC:12785] | -4.422 | -0.0673 | Yes |
| 50 | HHIP | hedgehog interacting protein [Source:HGNC Symbol;Acc:HGNC:14866] | -4.454 | 0.0019 | Yes |

| Oxidative phosphorylation | SYMBOL | TITLE | RANK METRIC SCORE | RUNNING ES | CORE ENRICHMENT |
|---|---|---|---|---|---|
| 1 | ATP6V0D2 | ATPase H+ transporting V0 subunit d2 [Source:HGNC Symbol;Acc:HGNC:18266] | 1.659 | -0.0338 | Yes |
| 2 | COX6A2 | cytochrome c oxidase subunit 6A2 [Source:HGNC Symbol;Acc:HGNC:2279] | 1.463 | -0.0294 | Yes |
| 3 | ATP6V0A2 | ATPase H+ transporting V0 subunit a2 [Source:HGNC Symbol;Acc:HGNC:18481] | 1.25 | -0.0335 | Yes |
| 4 | NDUFS1 | NADH:ubiquinone oxidoreductase core subunit S1 [Source:HGNC Symbol;Acc:HGNC:7707] | 1.056 | -0.0457 | Yes |
| 5 | NDUFV1 | NADH:ubiquinone oxidoreductase core subunit V1 [Source:HGNC Symbol;Acc:HGNC:7716] | 1.006 | -0.0414 | Yes |
| 6 | UQCR10 | "ubiquinol-cytochrome c reductase, complex III subunit X [Source:HGNC Symbol;Acc:HGNC:30863]" | 0.974 | -0.0352 | Yes |
| 7 | COX4I1 | cytochrome c oxidase subunit 4I1 [Source:HGNC Symbol;Acc:HGNC:2265] | 0.968 | -0.0238 | Yes |
| 8 | ATP6V0A4 | ATPase H+ transporting V0 subunit a4 [Source:HGNC Symbol;Acc:HGNC:866] | 0.955 | -0.014 | Yes |
| 9 | MT-ND4 | mitochondrially encoded NADH:ubiquinone oxidoreductase core subunit 4 [Source:HGNC Symbol;Acc:HGNC:7459] | 0.932 | -0.0072 | Yes |
| 10 | SDHA | succinate dehydrogenase complex flavoprotein subunit A [Source:HGNC | 0.923 | 0.0027 | Yes |

| | | | | | |
|---|---|---|---|---|---|
| | | Symbol;Acc:HGNC:10680] | | | |
| 11 | ATP6AP1 | ATPase H+ transporting accessory protein 1 [Source:HGNC Symbol;Acc:HGNC:868] | 0.92 | 0.0137 | Yes |
| 12 | NDUFB10 | NADH:ubiquinone oxidoreductase subunit B10 [Source:HGNC Symbol;Acc:HGNC:7696] | 0.889 | 0.0185 | Yes |
| 13 | UQCR11 | "ubiquinol-cytochrome c reductase, complex III subunit XI [Source:HGNC Symbol;Acc:HGNC:30862]" | 0.883 | 0.0286 | Yes |
| 14 | NDUFA4 | NDUFA4 mitochondrial complex associated [Source:HGNC Symbol;Acc:HGNC:7687] | 0.86 | 0.0329 | Yes |
| 15 | ATP5MC1 | ATP synthase membrane subunit c locus 1 [Source:HGNC Symbol;Acc:HGNC:841] | 0.851 | 0.0412 | Yes |
| 16 | MT-ND2 | mitochondrially encoded NADH:ubiquinone oxidoreductase core subunit 2 [Source:HGNC Symbol;Acc:HGNC:7456] | 0.848 | 0.0514 | Yes |
| 17 | CYC1 | cytochrome c1 [Source:HGNC Symbol;Acc:HGNC:2579] | 0.845 | 0.0615 | Yes |
| 18 | UQCRQ | ubiquinol-cytochrome c reductase complex III subunit VII [Source:HGNC Symbol;Acc:HGNC:29594] | 0.842 | 0.0712 | Yes |
| 19 | MT-ATP6 | mitochondrially encoded ATP synthase membrane subunit 6 [Source:HGNC Symbol;Acc:HGNC:7414] | 0.838 | 0.0807 | Yes |
| 20 | COX5A | cytochrome c oxidase subunit 5A [Source:HGNC Symbol;Acc:HGNC:2267] | 0.836 | 0.0909 | Yes |
| 21 | ATP5MG | ATP synthase membrane subunit g [Source:HGNC Symbol;Acc:HGNC:14247] | 0.826 | 0.0987 | Yes |

| | | | | | |
|---|---|---|---|---|---|
| 22 | COX6B1 | cytochrome c oxidase subunit 6B1 [Source:HGNC Symbol;Acc:HGNC:2280] | 0.825 | 0.109 | Yes |
| 23 | ATP6V1F | ATPase H+ transporting V1 subunit F [Source:HGNC Symbol;Acc:HGNC:16832] | 0.81 | 0.115 | Yes |
| 24 | MT-ND4L | mitochondrially encoded NADH:ubiquinone oxidoreductase core subunit 4L [Source:HGNC Symbol;Acc:HGNC:7460] | 0.81 | 0.1253 | Yes |
| 25 | UQCRH | ubiquinol-cytochrome c reductase hinge protein [Source:HGNC Symbol;Acc:HGNC:12590] | 0.804 | 0.1337 | Yes |
| 26 | ATP6V0B | ATPase H+ transporting V0 subunit b [Source:HGNC Symbol;Acc:HGNC:861] | 0.802 | 0.1431 | Yes |
| 27 | NDUFA8 | NADH:ubiquinone oxidoreductase subunit A8 [Source:HGNC Symbol;Acc:HGNC:7692] | 0.8 | 0.1532 | Yes |
| 28 | NDUFA11 | NADH:ubiquinone oxidoreductase subunit A11 [Source:HGNC Symbol;Acc:HGNC:20371] | 0.798 | 0.1628 | Yes |
| 29 | ATP5MC3 | ATP synthase membrane subunit c locus 3 [Source:HGNC Symbol;Acc:HGNC:843] | 0.795 | 0.1721 | Yes |
| 30 | MT-ND1 | mitochondrially encoded NADH:ubiquinone oxidoreductase core subunit 1 [Source:HGNC Symbol;Acc:HGNC:7455] | 0.792 | 0.181 | Yes |
| 31 | COX7B | cytochrome c oxidase subunit 7B [Source:HGNC Symbol;Acc:HGNC:2291] | 0.781 | 0.1877 | Yes |
| 32 | COX8A | cytochrome c oxidase subunit 8A [Source:HGNC Symbol;Acc:HGNC:2294] | 0.755 | 0.1893 | Yes |

| | | | | | |
|---|---|---|---|---|---|
| 33 | COX7A2 | cytochrome c oxidase subunit 7A2 [Source:HGNC Symbol;Acc:HGNC:2288] | 0.747 | 0.1969 | Yes |
| 34 | MT-ND3 | mitochondrially encoded NADH:ubiquinone oxidoreductase core subunit 3 [Source:HGNC Symbol;Acc:HGNC:7458] | 0.742 | 0.2043 | Yes |
| 35 | NDUFAB1 | NADH:ubiquinone oxidoreductase subunit AB1 [Source:HGNC Symbol;Acc:HGNC:7694] | 0.741 | 0.2134 | Yes |
| 36 | ATP5PB | ATP synthase peripheral stalk-membrane subunit b [Source:HGNC Symbol;Acc:HGNC:840] | 0.736 | 0.2209 | Yes |
| 37 | NDUFC2 | NADH:ubiquinone oxidoreductase subunit C2 [Source:HGNC Symbol;Acc:HGNC:7706] | 0.729 | 0.2277 | Yes |
| 38 | ATP5F1B | ATP synthase F1 subunit beta [Source:HGNC Symbol;Acc:HGNC:830] | 0.727 | 0.2365 | Yes |
| 39 | COX11 | cytochrome c oxidase copper chaperone COX11 [Source:HGNC Symbol;Acc:HGNC:2261] | 0.72 | 0.2424 | Yes |
| 40 | NDUFA9 | NADH:ubiquinone oxidoreductase subunit A9 [Source:HGNC Symbol;Acc:HGNC:7693] | 0.718 | 0.2511 | Yes |
| 41 | ATP5F1D | ATP synthase F1 subunit delta [Source:HGNC Symbol;Acc:HGNC:837] | 0.707 | 0.2564 | Yes |
| 42 | NDUFB9 | NADH:ubiquinone oxidoreductase subunit B9 [Source:HGNC Symbol;Acc:HGNC:7704] | 0.706 | 0.2651 | Yes |
| 43 | ATP5PD | ATP synthase peripheral stalk subunit d [Source:HGNC Symbol;Acc:HGNC:845] | 0.702 | 0.2725 | Yes |
| 44 | MT-CYB | mitochondrially encoded cytochrome b [Source:HGNC Symbol;Acc:HGNC:7427] | 0.693 | 0.2777 | Yes |
| 45 | MT-ATP8 | mitochondrially encoded ATP synthase membrane | 0.687 | 0.2841 | Yes |

| | | | | | |
|---|---|---|---|---|---|
| | | subunit 8 [Source:HGNC Symbol;Acc:HGNC:7415] | | | |
| 46 | NDUFB6 | NADH:ubiquinone oxidoreductase subunit B6 [Source:HGNC Symbol;Acc:HGNC:7701] | 0.687 | 0.2929 | Yes |
| 47 | ATP5ME | ATP synthase membrane subunit e [Source:HGNC Symbol;Acc:HGNC:846] | 0.687 | 0.3015 | Yes |
| 48 | MT-CO3 | mitochondrially encoded cytochrome c oxidase III [Source:HGNC Symbol;Acc:HGNC:7422] | 0.685 | 0.31 | Yes |
| 49 | UQCRB | ubiquinol-cytochrome c reductase binding protein [Source:HGNC Symbol;Acc:HGNC:1258 2] | 0.673 | 0.3134 | Yes |
| 50 | SDHC | succinate dehydrogenase complex subunit C [Source:HGNC Symbol;Acc:HGNC:1068 2] | 0.668 | 0.3197 | Yes |
| 51 | MT-CO1 | mitochondrially encoded cytochrome c oxidase I [Source:HGNC Symbol;Acc:HGNC:7419] | 0.665 | 0.3271 | Yes |
| 52 | COX6A1 | cytochrome c oxidase subunit 6A1 [Source:HGNC Symbol;Acc:HGNC:2277] | 0.664 | 0.3353 | Yes |
| 53 | NDUFS5 | NADH:ubiquinone oxidoreductase subunit S5 [Source:HGNC Symbol;Acc:HGNC:7712] | 0.663 | 0.3435 | Yes |
| 54 | COX15 | cytochrome c oxidase assembly homolog COX15 [Source:HGNC Symbol;Acc:HGNC:2263] | 0.66 | 0.3503 | Yes |
| 55 | NDUFA10 | NADH:ubiquinone oxidoreductase subunit A10 [Source:HGNC Symbol;Acc:HGNC:7684] | 0.646 | 0.3532 | Yes |
| 56 | SDHD | succinate dehydrogenase complex subunit D [Source:HGNC Symbol;Acc:HGNC:1068 3] | 0.645 | 0.3611 | Yes |
| 57 | NDUFB7 | NADH:ubiquinone oxidoreductase subunit B7 | 0.643 | 0.3686 | Yes |

| | | | | | |
|---|---|---|---|---|---|
| | | [Source:HGNC Symbol;Acc:HGNC:7702] | | | |
| 58 | SDHB | succinate dehydrogenase complex iron sulfur subunit B [Source:HGNC Symbol;Acc:HGNC:10681] | 0.639 | 0.3751 | Yes |
| 59 | NDUFB5 | NADH:ubiquinone oxidoreductase subunit B5 [Source:HGNC Symbol;Acc:HGNC:7700] | 0.639 | 0.3829 | Yes |
| 60 | UQCRFS1 | "ubiquinol-cytochrome c reductase, Rieske iron-sulfur polypeptide 1 [Source:HGNC Symbol;Acc:HGNC:12587]" | 0.637 | 0.3902 | Yes |
| 61 | COX5B | cytochrome c oxidase subunit 5B [Source:HGNC Symbol;Acc:HGNC:2269] | 0.634 | 0.3971 | Yes |
| 62 | COX10 | cytochrome c oxidase assembly factor heme A:farnesyltransferase COX10 [Source:HGNC Symbol;Acc:HGNC:2260] | 0.634 | 0.405 | Yes |
| 63 | UQCRC2 | ubiquinol-cytochrome c reductase core protein 2 [Source:HGNC Symbol;Acc:HGNC:12586] | 0.618 | 0.4038 | Yes |
| 64 | ATP6V1C1 | ATPase H+ transporting V1 subunit C1 [Source:HGNC Symbol;Acc:HGNC:856] | 0.614 | 0.4096 | Yes |
| 65 | COX7C | cytochrome c oxidase subunit 7C [Source:HGNC Symbol;Acc:HGNC:2292] | 0.611 | 0.4161 | Yes |
| 66 | MT-ND5 | mitochondrially encoded NADH:ubiquinone oxidoreductase core subunit 5 [Source:HGNC Symbol;Acc:HGNC:7461] | 0.611 | 0.4237 | Yes |
| 67 | NDUFS7 | NADH:ubiquinone oxidoreductase core subunit S7 [Source:HGNC Symbol;Acc:HGNC:7714] | 0.605 | 0.4288 | Yes |
| 68 | ATP5PF | ATP synthase peripheral stalk subunit F6 | 0.604 | 0.4358 | Yes |

| | | | | | |
|---|---|---|---|---|---|
| | | [Source:HGNC Symbol;Acc:HGNC:847] | | | |
| 69 | NDUFV2 | NADH:ubiquinone oxidoreductase core subunit V2 [Source:HGNC Symbol;Acc:HGNC:7717] | 0.601 | 0.4424 | Yes |
| 70 | NDUFS3 | NADH:ubiquinone oxidoreductase core subunit S3 [Source:HGNC Symbol;Acc:HGNC:7710] | 0.597 | 0.448 | Yes |
| 71 | ATP6V1B2 | ATPase H+ transporting V1 subunit B2 [Source:HGNC Symbol;Acc:HGNC:854] | 0.596 | 0.4551 | Yes |
| 72 | ATP6V1A | ATPase H+ transporting V1 subunit A [Source:HGNC Symbol;Acc:HGNC:851] | 0.589 | 0.4581 | Yes |
| 73 | UQCRC1 | ubiquinol-cytochrome c reductase core protein 1 [Source:HGNC Symbol;Acc:HGNC:12585] | 0.576 | 0.4595 | Yes |
| 74 | NDUFA7 | NADH:ubiquinone oxidoreductase subunit A7 [Source:HGNC Symbol;Acc:HGNC:7691] | 0.561 | 0.4582 | Yes |
| 75 | COX7A2L | cytochrome c oxidase subunit 7A2 like [Source:HGNC Symbol;Acc:HGNC:2289] | 0.55 | 0.4601 | Yes |
| 76 | ATP5MF | ATP synthase membrane subunit f [Source:HGNC Symbol;Acc:HGNC:848] | 0.55 | 0.467 | Yes |
| 77 | ATP6V0E1 | ATPase H+ transporting V0 subunit e1 [Source:HGNC Symbol;Acc:HGNC:863] | 0.549 | 0.4739 | Yes |
| 78 | ATP5F1C | ATP synthase F1 subunit gamma [Source:HGNC Symbol;Acc:HGNC:833] | 0.536 | 0.4734 | Yes |
| 79 | NDUFC1 | NADH:ubiquinone oxidoreductase subunit C1 [Source:HGNC Symbol;Acc:HGNC:7705] | 0.527 | 0.4749 | Yes |
| 80 | NDUFB2 | NADH:ubiquinone oxidoreductase subunit B2 [Source:HGNC Symbol;Acc:HGNC:7697] | 0.523 | 0.4792 | Yes |

| | | | | | |
|---|---|---|---|---|---|
| 81 | COX6C | cytochrome c oxidase subunit 6C [Source:HGNC Symbol;Acc:HGNC:2285] | 0.509 | 0.4766 | Yes |
| 82 | NDUFS6 | NADH:ubiquinone oxidoreductase subunit S6 [Source:HGNC Symbol;Acc:HGNC:7713] | 0.505 | 0.4803 | Yes |
| 83 | ATP6V1H | ATPase H+ transporting V1 subunit H [Source:HGNC Symbol;Acc:HGNC:18303] | 0.504 | 0.4859 | Yes |
| 84 | NDUFA5 | NADH:ubiquinone oxidoreductase subunit A5 [Source:HGNC Symbol;Acc:HGNC:7688] | 0.5 | 0.4896 | Yes |
| 85 | NDUFS2 | NADH:ubiquinone oxidoreductase core subunit S2 [Source:HGNC Symbol;Acc:HGNC:7708] | 0.5 | 0.4951 | Yes |

| Citrate and TCA cycle | SYMBOL | TITLE | RANK METRIC SCORE | RUNNING ES | CORE ENRICHMENT |
|---|---|---|---|---|---|
| 1 | PC | pyruvate carboxylase [Source:HGNC Symbol;Acc:HGNC:8636] | 1.458 | 0.0068 | Yes |
| 2 | IDH1 | isocitrate dehydrogenase (NADP(+)) 1 [Source:HGNC Symbol;Acc:HGNC:5382] | 1.069 | 0.0202 | Yes |
| 3 | IDH3A | isocitrate dehydrogenase (NAD(+)) 3 catalytic subunit alpha [Source:HGNC Symbol;Acc:HGNC:5384] | 1.047 | 0.0707 | Yes |
| 4 | SDHA | succinate dehydrogenase complex flavoprotein subunit A [Source:HGNC Symbol;Acc:HGNC:10680] | 0.923 | 0.0956 | Yes |
| 5 | SUCLG2 | succinate-CoA ligase GDP-forming subunit beta [Source:HGNC Symbol;Acc:HGNC:11450] | 0.923 | 0.1439 | Yes |
| 6 | MDH1 | malate dehydrogenase 1 [Source:HGNC Symbol;Acc:HGNC:6970] | 0.897 | 0.1858 | Yes |
| 7 | ACO2 | aconitase 2 [Source:HGNC Symbol;Acc:HGNC:118] | 0.892 | 0.2314 | Yes |

| | | | | | |
|---|---|---|---|---|---|
| 8 | OGDH | oxoglutarate dehydrogenase [Source:HGNC Symbol;Acc:HGNC:8124] | 0.884 | 0.2761 | Yes |
| 9 | IDH2 | isocitrate dehydrogenase (NADP(+)) 2 [Source:HGNC Symbol;Acc:HGNC:5383] | 0.837 | 0.306 | Yes |
| 10 | SUCLG1 | succinate-CoA ligase GDP/ADP-forming subunit alpha [Source:HGNC Symbol;Acc:HGNC:11449] | 0.796 | 0.3356 | Yes |
| 11 | MDH2 | malate dehydrogenase 2 [Source:HGNC Symbol;Acc:HGNC:6971] | 0.794 | 0.3763 | Yes |
| 12 | FH | fumarate hydratase [Source:HGNC Symbol;Acc:HGNC:3700] | 0.761 | 0.4062 | Yes |
| 13 | DLD | dihydrolipoamide dehydrogenase [Source:HGNC Symbol;Acc:HGNC:2898] | 0.734 | 0.4351 | Yes |
| 14 | IDH3G | isocitrate dehydrogenase (NAD(+)) 3 non-catalytic subunit gamma [Source:HGNC Symbol;Acc:HGNC:5386] | 0.715 | 0.4654 | Yes |
| 15 | DLST | dihydrolipoamide S-succinyltransferase [Source:HGNC Symbol;Acc:HGNC:2911] | 0.7 | 0.4961 | Yes |
| 16 | SDHC | succinate dehydrogenase complex subunit C [Source:HGNC Symbol;Acc:HGNC:10682] | 0.668 | 0.5179 | Yes |
| 17 | SDHD | succinate dehydrogenase complex subunit D [Source:HGNC Symbol;Acc:HGNC:10683] | 0.645 | 0.5423 | Yes |
| 18 | SDHB | succinate dehydrogenase complex iron sulfur subunit B [Source:HGNC Symbol;Acc:HGNC:10681] | 0.639 | 0.5733 | Yes |
| 19 | SUCLA2 | succinate-CoA ligase ADP-forming subunit beta [Source:HGNC Symbol;Acc:HGNC:11448] | 0.539 | 0.5478 | Yes |
| 20 | PDHB | pyruvate dehydrogenase E1 subunit beta [Source:HGNC Symbol;Acc:HGNC:8808] | 0.526 | 0.5684 | Yes |
| 21 | DLAT | dihydrolipoamide S-acetyltransferase [Source:HGNC Symbol;Acc:HGNC:2896] | 0.5 | 0.5761 | Yes |

| | | | | | |
|---|---|---|---|---|---|
| 22 | CS | citrate synthase [Source:HGNC Symbol;Acc:HGNC:2422] | 0.463 | 0.5749 | Yes |
| 23 | IDH3B | isocitrate dehydrogenase (NAD(+)) 3 non-catalytic subunit beta [Source:HGNC Symbol;Acc:HGNC:5385] | 0.437 | 0.5798 | Yes |
| 24 | PDHA1 | pyruvate dehydrogenase E1 subunit alpha 1 [Source:HGNC Symbol;Acc:HGNC:8806] | 0.415 | 0.5861 | Yes |
| 25 | ACLY | ATP citrate lyase [Source:HGNC Symbol;Acc:HGNC:115] | 0.413 | 0.6061 | Yes |

| Porphyrin and chlorophyll metabolism | SYMBOL | TITLE | RANK METRIC SCORE | RUNNING ES | CORE ENRICHMENT |
|---|---|---|---|---|---|
| 1 | UGT1A4 | UDP glucuronosyltransferase family 1 member A4 [Source:HGNC Symbol;Acc:HGNC:12536] | 3.083 | 0.1297 | Yes |
| 2 | UGT1A6 | UDP glucuronosyltransferase family 1 member A6 [Source:HGNC Symbol;Acc:HGNC:12538] | 2.674 | 0.2454 | Yes |
| 3 | UGT1A3 | UDP glucuronosyltransferase family 1 member A3 [Source:HGNC Symbol;Acc:HGNC:12535] | 2.61 | 0.3638 | Yes |
| 4 | UGT1A10 | UDP glucuronosyltransferase family 1 member A10 [Source:HGNC Symbol;Acc:HGNC:12531] | 1.855 | 0.4241 | Yes |
| 5 | HMOX1 | heme oxygenase 1 [Source:HGNC Symbol;Acc:HGNC:5013] | 1.604 | 0.4854 | Yes |
| 6 | FTH1 | ferritin heavy chain 1 [Source:HGNC Symbol;Acc:HGNC:3976] | 1.166 | 0.4975 | Yes |
| 7 | ALAS1 | 5'-aminolevulinate synthase 1 [Source:HGNC Symbol;Acc:HGNC:396] | 0.929 | 0.5007 | Yes |
| 8 | BLVRB | biliverdin reductase B [Source:HGNC Symbol;Acc:HGNC:1063] | 0.907 | 0.5379 | Yes |
| 9 | FECH | ferrochelatase [Source:HGNC Symbol;Acc:HGNC:3647] | 0.833 | 0.5558 | Yes |

| | | | | | |
|---|---|---|---|---|---|
| 10 | UROS | uroporphyrinogen III synthase [Source:HGNC Symbol;Acc:HGNC:12592] | 0.664 | 0.5258 | Yes |
| 11 | COX15 | cytochrome c oxidase assembly homolog COX15 [Source:HGNC Symbol;Acc:HGNC:2263] | 0.66 | 0.5543 | Yes |
| 12 | COX10 | cytochrome c oxidase assembly factor heme A:farnesyltransferase COX10 [Source:HGNC Symbol;Acc:HGNC:2260] | 0.634 | 0.5721 | Yes |
| 13 | GUSB | glucuronidase beta [Source:HGNC Symbol;Acc:HGNC:4696] | 0.633 | 0.6007 | Yes |
| 14 | EPRS1 | glutamyl-prolyl-tRNA synthetase 1 [Source:HGNC Symbol;Acc:HGNC:3418] | 0.595 | 0.6073 | Yes |
| 15 | BLVRA | biliverdin reductase A [Source:HGNC Symbol;Acc:HGNC:1062] | 0.58 | 0.6259 | Yes |

| Primary immunodeficiency | SYMBOL | TITLE | RANK METRIC SCORE | RUNNING ES | CORE ENRICHMENT |
|---|---|---|---|---|---|
| 15 | CD79A | CD79a molecule [Source:HGNC Symbol;Acc:HGNC:1698] | -0.637 | -0.5689 | Yes |
| 16 | JAK3 | Janus kinase 3 [Source:HGNC Symbol;Acc:HGNC:6193] | -0.671 | -0.5407 | Yes |
| 17 | TNFRSF13C | TNF receptor superfamily member 13C [Source:HGNC Symbol;Acc:HGNC:17755] | -0.917 | -0.5225 | Yes |
| 18 | CIITA | class II major histocompatibility complex transactivator [Source:HGNC Symbol;Acc:HGNC:7067] | -1.181 | -0.4867 | Yes |
| 19 | UNG | uracil DNA glycosylase [Source:HGNC Symbol;Acc:HGNC:12572] | -1.446 | -0.4357 | Yes |
| 20 | ZAP70 | zeta chain of T cell receptor associated protein kinase 70 [Source:HGNC Symbol;Acc:HGNC:12858] | -1.983 | -0.3661 | Yes |
| 21 | LCK | "LCK proto-oncogene, Src family tyrosine kinase [Source:HGNC Symbol;Acc:HGNC:6524]" | -2.064 | -0.2717 | Yes |

| | | | | | |
|---|---|---|---|---|---|
| 22 | BLNK | B cell linker [Source:HGNC Symbol;Acc:HGNC:14211] | -3.143 | -0.1565 | Yes |
| 23 | CD40 | CD40 molecule [Source:HGNC Symbol;Acc:HGNC:11919] | -3.724 | 0.0096 | Yes |

| Val Leu and Ile (BCAA) degradation | SYMBOL | TITLE | RANK METRIC SCORE | RUNNING ES | CORE ENRICHMENT |
|---|---|---|---|---|---|
| 1 | HMGCS2 | 3-hydroxy-3-methylglutaryl-CoA synthase 2 [Source:HGNC Symbol;Acc:HGNC:5008] | 1.921 | 0.0204 | Yes |
| 2 | BCAT1 | branched chain amino acid transaminase 1 [Source:HGNC Symbol;Acc:HGNC:976] | 1.312 | 0.0246 | Yes |
| 3 | ABAT | 4-aminobutyrate aminotransferase [Source:HGNC Symbol;Acc:HGNC:23] | 1.198 | 0.0516 | Yes |
| 4 | AUH | AU RNA binding methylglutaconyl-CoA hydratase [Source:HGNC Symbol;Acc:HGNC:890] | 1.196 | 0.0906 | Yes |
| 5 | ALDH6A1 | aldehyde dehydrogenase 6 family member A1 [Source:HGNC Symbol;Acc:HGNC:7179] | 1.161 | 0.1241 | Yes |
| 6 | ACADSB | acyl-CoA dehydrogenase short/branched chain [Source:HGNC Symbol;Acc:HGNC:91] | 1.13 | 0.157 | Yes |
| 7 | MMUT | methylmalonyl-CoA mutase [Source:HGNC Symbol;Acc:HGNC:7526] | 1.129 | 0.1942 | Yes |
| 8 | ALDH3A2 | aldehyde dehydrogenase 3 family member A2 [Source:HGNC Symbol;Acc:HGNC:403] | 1.048 | 0.2171 | Yes |
| 9 | HADHB | hydroxyacyl-CoA dehydrogenase trifunctional multienzyme complex subunit beta [Source:HGNC Symbol;Acc:HGNC:4803] | 1.032 | 0.2482 | Yes |
| 10 | ACAA2 | acetyl-CoA acyltransferase 2 [Source:HGNC Symbol;Acc:HGNC:83] | 0.989 | 0.2737 | Yes |

| | | | | | |
|---|---|---|---|---|---|
| 11 | BCKDHB | branched chain keto acid dehydrogenase E1 subunit beta [Source:HGNC Symbol;Acc:HGNC:987] | 0.968 | 0.3014 | Yes |
| 12 | ALDH1B1 | aldehyde dehydrogenase 1 family member B1 [Source:HGNC Symbol;Acc:HGNC:407] | 0.936 | 0.3257 | Yes |
| 13 | BCKDHA | branched chain keto acid dehydrogenase E1 subunit alpha [Source:HGNC Symbol;Acc:HGNC:986] | 0.91 | 0.3504 | Yes |
| 14 | HMGCS1 | 3-hydroxy-3-methylglutaryl-CoA synthase 1 [Source:HGNC Symbol;Acc:HGNC:5007] | 0.825 | 0.3543 | Yes |
| 15 | ACADM | acyl-CoA dehydrogenase medium chain [Source:HGNC Symbol;Acc:HGNC:89] | 0.784 | 0.3667 | Yes |
| 16 | PCCB | propionyl-CoA carboxylase subunit beta [Source:HGNC Symbol;Acc:HGNC:8654] | 0.751 | 0.382 | Yes |
| 17 | ACAA1 | acetyl-CoA acyltransferase 1 [Source:HGNC Symbol;Acc:HGNC:82] | 0.743 | 0.4034 | Yes |
| 18 | HIBADH | 3-hydroxyisobutyrate dehydrogenase [Source:HGNC Symbol;Acc:HGNC:4907] | 0.737 | 0.4257 | Yes |
| 19 | DLD | dihydrolipoamide dehydrogenase [Source:HGNC Symbol;Acc:HGNC:2898] | 0.734 | 0.4486 | Yes |
| 20 | ACAT1 | acetyl-CoA acetyltransferase 1 [Source:HGNC Symbol;Acc:HGNC:93] | 0.727 | 0.4702 | Yes |
| 21 | PCCA | propionyl-CoA carboxylase subunit alpha [Source:HGNC Symbol;Acc:HGNC:8653] | 0.704 | 0.485 | Yes |
| 22 | IVD | isovaleryl-CoA dehydrogenase [Source:HGNC Symbol;Acc:HGNC:6186] | 0.692 | 0.5029 | Yes |
| 23 | OXCT1 | 3-oxoacid CoA-transferase 1 [Source:HGNC Symbol;Acc:HGNC:8527] | 0.632 | 0.4975 | Yes |

| | | | | | |
|---|---|---|---|---|---|
| 24 | MCEE | methylmalonyl-CoA epimerase [Source:HGNC Symbol;Acc:HGNC:16732] | 0.618 | 0.5098 | Yes |
| 25 | BCAT2 | branched chain amino acid transaminase 2 [Source:HGNC Symbol;Acc:HGNC:977] | 0.573 | 0.5046 | Yes |
| 26 | HADHA | hydroxyacyl-CoA dehydrogenase trifunctional multienzyme complex subunit alpha [Source:HGNC Symbol;Acc:HGNC:4801] | 0.538 | 0.5034 | Yes |
| 27 | MCCC2 | methylcrotonoyl-CoA carboxylase 2 [Source:HGNC Symbol;Acc:HGNC:6937] | 0.511 | 0.5043 | Yes |
| 28 | ALDH7A1 | aldehyde dehydrogenase 7 family member A1 [Source:HGNC Symbol;Acc:HGNC:877] | 0.477 | 0.4949 | Yes |
| 29 | HIBCH | 3-hydroxyisobutyryl-CoA hydrolase [Source:HGNC Symbol;Acc:HGNC:4908] | 0.468 | 0.5048 | Yes |
| 30 | MCCC1 | methylcrotonoyl-CoA carboxylase 1 [Source:HGNC Symbol;Acc:HGNC:6936] | 0.462 | 0.5156 | Yes |
| 31 | HADH | hydroxyacyl-CoA dehydrogenase [Source:HGNC Symbol;Acc:HGNC:4799] | 0.456 | 0.5259 | Yes |

| Cell cycle | SYMBOL | TITLE | RANK METRIC SCORE | RUNNING ES | CORE ENRICHMENT |
|---|---|---|---|---|---|
| 89 | DBF4 | DBF4 zinc finger [Source:HGNC Symbol;Acc:HGNC:17364] | -0.475 | -0.4259 | Yes |
| 90 | CDC20 | cell division cycle 20 [Source:HGNC Symbol;Acc:HGNC:1723] | -0.511 | -0.4243 | Yes |
| 91 | CDK1 | cyclin dependent kinase 1 [Source:HGNC Symbol;Acc:HGNC:1722] | -0.563 | -0.4237 | Yes |
| 92 | SKP2 | S-phase kinase associated protein 2 [Source:HGNC Symbol;Acc:HGNC:10901] | -0.632 | -0.4256 | Yes |

| | | | | | |
|---|---|---|---|---|---|
| 93 | CDC25B | cell division cycle 25B [Source:HGNC Symbol;Acc:HGNC:1726] | -0.648 | -0.4186 | Yes |
| 94 | CHEK1 | checkpoint kinase 1 [Source:HGNC Symbol;Acc:HGNC:1925] | -0.724 | -0.4165 | Yes |
| 95 | MAD2L1 | mitotic arrest deficient 2 like 1 [Source:HGNC Symbol;Acc:HGNC:6763] | -0.74 | -0.4085 | Yes |
| 96 | MCM6 | minichromosome maintenance complex component 6 [Source:HGNC Symbol;Acc:HGNC:6949] | -0.755 | -0.4005 | Yes |
| 97 | CDK6 | cyclin dependent kinase 6 [Source:HGNC Symbol;Acc:HGNC:1777] | -0.77 | -0.3921 | Yes |
| 98 | GADD45A | growth arrest and DNA damage inducible alpha [Source:HGNC Symbol;Acc:HGNC:4095] | -0.812 | -0.3856 | Yes |
| 99 | CCND2 | cyclin D2 [Source:HGNC Symbol;Acc:HGNC:1583] | -0.812 | -0.3748 | Yes |
| 100 | CDC25A | cell division cycle 25A [Source:HGNC Symbol;Acc:HGNC:1725] | -0.842 | -0.3665 | Yes |
| 101 | CDC14A | cell division cycle 14A [Source:HGNC Symbol;Acc:HGNC:1718] | -0.876 | -0.3574 | Yes |
| 102 | CHEK2 | checkpoint kinase 2 [Source:HGNC Symbol;Acc:HGNC:16627] | -0.901 | -0.3478 | Yes |
| 103 | ORC1 | origin recognition complex subunit 1 [Source:HGNC Symbol;Acc:HGNC:8487] | -0.979 | -0.3413 | Yes |
| 104 | CCNE1 | cyclin E1 [Source:HGNC Symbol;Acc:HGNC:1589] | -1.112 | -0.3367 | Yes |
| 105 | TGFB3 | transforming growth factor beta 3 [Source:HGNC Symbol;Acc:HGNC:11769] | -1.196 | -0.3269 | Yes |
| 106 | CCNE2 | cyclin E2 [Source:HGNC Symbol;Acc:HGNC:1590] | -1.36 | -0.3195 | Yes |
| 107 | BUB1B | BUB1 mitotic checkpoint serine/threonine kinase B [Source:HGNC Symbol;Acc:HGNC:1149] | -1.416 | -0.3037 | Yes |
| 108 | MCM3 | minichromosome maintenance complex component 3 | -1.446 | -0.2866 | Yes |

| | | | | | |
|---|---|---|---|---|---|
| | | [Source:HGNC Symbol;Acc:HGNC:6945] | | | |
| 109 | MCM2 | minichromosome maintenance complex component 2 [Source:HGNC Symbol;Acc:HGNC:6944] | -1.486 | -0.2689 | Yes |
| 110 | MCM5 | minichromosome maintenance complex component 5 [Source:HGNC Symbol;Acc:HGNC:6948] | -1.588 | -0.2527 | Yes |
| 111 | CCNA2 | cyclin A2 [Source:HGNC Symbol;Acc:HGNC:1578] | -1.71 | -0.2351 | Yes |
| 112 | PLK1 | polo like kinase 1 [Source:HGNC Symbol;Acc:HGNC:9077] | -1.884 | -0.2172 | Yes |
| 113 | CCND1 | cyclin D1 [Source:HGNC Symbol;Acc:HGNC:1582] | -1.886 | -0.192 | Yes |
| 114 | TTK | TTK protein kinase [Source:HGNC Symbol;Acc:HGNC:12401] | -1.918 | -0.1676 | Yes |
| 115 | BUB1 | BUB1 mitotic checkpoint serine/threonine kinase [Source:HGNC Symbol;Acc:HGNC:1148] | -1.99 | -0.1447 | Yes |
| 116 | CDC25C | cell division cycle 25C [Source:HGNC Symbol;Acc:HGNC:1727] | -2.062 | -0.1201 | Yes |
| 117 | CDC6 | cell division cycle 6 [Source:HGNC Symbol;Acc:HGNC:1744] | -2.36 | -0.1002 | Yes |
| 118 | SMC1B | structural maintenance of chromosomes 1B [Source:HGNC Symbol;Acc:HGNC:11112] | -2.363 | -0.0688 | Yes |
| 119 | CCNB2 | cyclin B2 [Source:HGNC Symbol;Acc:HGNC:1580] | -2.515 | -0.0406 | Yes |
| 120 | E2F2 | E2F transcription factor 2 [Source:HGNC Symbol;Acc:HGNC:3114] | -2.62 | -0.0083 | Yes |
| 121 | CCNA1 | cyclin A1 [Source:HGNC Symbol;Acc:HGNC:1577] | -3.162 | 0.0194 | Yes |

| **Drug metabolism** | **SYMBOL** | **TITLE** | **RANK METRIC SCORE** | **RUNNING ES** | **CORE ENRICHMENT** |
|---|---|---|---|---|---|

| | | | | | |
|---|---|---|---|---|---|
| 1 | XDH | xanthine dehydrogenase [Source:HGNC Symbol;Acc:HGNC:12805] | 3.167 | 0.0998 | Yes |
| 2 | UGT1A4 | UDP glucuronosyltransferase family 1 member A4 [Source:HGNC Symbol;Acc:HGNC:12536] | 3.083 | 0.2068 | Yes |
| 3 | UGT1A6 | UDP glucuronosyltransferase family 1 member A6 [Source:HGNC Symbol;Acc:HGNC:12538] | 2.674 | 0.2932 | Yes |
| 4 | DPYS | dihydropyrimidinase [Source:HGNC Symbol;Acc:HGNC:3013] | 2.641 | 0.3846 | Yes |
| 5 | UGT1A3 | UDP glucuronosyltransferase family 1 member A3 [Source:HGNC Symbol;Acc:HGNC:12535] | 2.61 | 0.4756 | Yes |
| 6 | CES5A | carboxylesterase 5A [Source:HGNC Symbol;Acc:HGNC:26459] | 1.972 | 0.524 | Yes |
| 7 | UGT1A10 | UDP glucuronosyltransferase family 1 member A10 [Source:HGNC Symbol;Acc:HGNC:12531] | 1.855 | 0.5846 | Yes |
| 8 | NAT2 | N-acetyltransferase 2 [Source:HGNC Symbol;Acc:HGNC:7646] | 1.631 | 0.631 | Yes |
| 9 | CES1 | carboxylesterase 1 [Source:HGNC Symbol;Acc:HGNC:1863] | 1.593 | 0.6845 | Yes |

| **Maturity onset diabetes of young** | **SYMBOL** | **TITLE** | **RANK METRIC SCORE** | **RUNNING ES** | **CORE ENRICHMENT** |
|---|---|---|---|---|---|
| 11 | NKX2-2 | NK2 homeobox 2 [Source:HGNC Symbol;Acc:HGNC:7835] | -1.472 | -0.5571 | Yes |
| 12 | HHEX | hematopoietically expressed homeobox [Source:HGNC Symbol;Acc:HGNC:4901] | -2.49 | -0.481 | Yes |

| | | | | | |
|---|---|---|---|---|---|
| 13 | MAFA | MAF bZIP transcription factor A [Source:HGNC Symbol;Acc:HGNC:23145] | -2.49 | -0.3623 | Yes |
| 14 | GCK | glucokinase [Source:HGNC Symbol;Acc:HGNC:4195] | -2.631 | -0.2408 | Yes |
| 15 | FOXA3 | forkhead box A3 [Source:HGNC Symbol;Acc:HGNC:5023] | -2.664 | -0.1143 | Yes |
| 16 | NKX6-1 | NK6 homeobox 1 [Source:HGNC Symbol;Acc:HGNC:7839] | -3.083 | 0.0214 | Yes |
